# Assessing the effects of warming and carbonate chemistry parameters on marine microbes in the Gulf of Mexico through basin-scale DNA metabarcoding

**DOI:** 10.1101/2024.07.30.605667

**Authors:** Sean R. Anderson, Katherine Silliman, Leticia Barbero, Fabian A. Gomez, Beth A. Stauffer, Astrid Schnetzer, Christopher R. Kelble, Luke R. Thompson

**Author notes:** National Oceanic and Atmospheric Administration, National Marine Fisheries Service, Office of Science and Technology (Christopher R. Kelble). Corresponding authors: Sean R. Anderson and Luke R. Thompson **Emails:** and.

## Abstract

Ocean acidification and warming threaten marine life, yet the impact of these processes on microbes remains unclear. Here, we performed basin-scale DNA metabarcoding of prokaryotes (16S V4–V5) and protists (18S V9) in the Gulf of Mexico and applied generalized linear models to reveal group-specific environmental correlates of functionally diverse microbes. Models supported prior physiological trends for some groups, like positive temperature effects on SAR11 and SAR86, and a positive effect of pH on *Prochlorococcus* that implied a negative response to decreasing pH. New insights were revealed for protists, like Syndiniales and Sagenista (e.g., positive pH effects), which offset positive relationships with temperature and reinforced the importance of considering multiple stressors simultaneously. Indicator analysis revealed phytoplankton, like *Ostreococcus* sp. and *Emiliania huxleyi*, that were associated with more acidic waters and may reflect candidate indicators of ocean change. Our findings highlight the need for sustained microbial sampling in marine systems, with implications for carbon export, nutrient cycling, and ecosystem health.

## Introduction

Our ability to predict how marine ecosystems and resources will respond to future ocean conditions will require accurate monitoring of marine biodiversity over space, time, and across natural environmental gradients (*1*). The oceans are changing rapidly, heavily impacted by rising concentrations of human-derived atmospheric carbon dioxide (CO_2_) that is absorbed at the ocean’s surface (*2*). Atmospheric CO_2_ has increased by nearly 50% (∼420 ppm at present) over the last century, leading to increased levels of dissolved inorganic carbon (DIC) in the ocean, in turn lowering seawater pH (*3*). This process of ocean acidification (OA) reduces saturation states for carbonate minerals, placing stress on organisms that require these minerals for cellular growth and other functions (*4*, *5*). The effects of OA are amplified by ocean warming, particularly at low latitudes, with surface temperatures expected to increase by 1–10 °C over the next century (*6*). Changes in seawater chemistry and physics can have immense impacts, both direct and indirect, on marine life (*3*). Thus, it is imperative to understand better how diverse marine organisms respond to present-day chemical and physical conditions to inform future potential shifts in community composition.

Over the past decade, research on species sensitivity to OA has expanded greatly, particularly for multicellular organisms that rely on carbonate chemistry for their structure and function (*3*, *5*). Much less research has been conducted on marine microbes (i.e., protists, Bacteria, and Archaea), despite the central role of microbes in food webs and their strong influence on biogeochemical cycles and carbon export (*7*, *8*). Microbes also respond quickly to shifts in their surrounding environment, making them potentially important indicators of changing ocean conditions (*9*, *10*). In general, global ecosystem models predict a decline in photosynthetic biomass and a shift in composition from larger plankton (e.g., diatoms) to picophytoplankton (0.2–2 µm), primarily driven by warming and enhanced stratification (*11–14*). Field and laboratory experiments have measured direct and negative impacts of OA on plankton, notably among calcifying haptophytes (e.g., coccolithophores), where increased partial pressure of CO_2_ (*p*CO_2_) and/or decreasing pH has led to reduced growth and calcification rates (*15–17*). However, evidence suggests that some phytoplankton species, even coccolithophores, may be resilient to rising *p*CO_2_ and warming (*13*, *18*, *19*). In addition, heterotrophic bacteria may be more resilient to OA compared to phytoplankton, impacted more directly by warming and changes to phytoplankton-derived organic matter (*20*, *21*). Employing DNA metabarcoding to characterize the complex effects of OA parameters and temperature on a wide range of microbes (*22*, *23*) will help guide lab-based experiments, identify indicator taxa, and inform model predictions.

The Gulf of Mexico (GOM) is an ideal location to study the effects of multiple stressors on marine microbes, as microbial communities in the GOM are affected by several major hydrographic features that result in strong physicochemical gradients (*24*). The GOM is a semi-enclosed subtropical basin, influenced by the Loop Current (and associated anticyclonic eddies) and freshwater inflow from riverine systems (Mississippi-Atchafalaya) in the north (*25*, *26*). Most of the GOM is oligotrophic (and nutrient-limited), with phytoplankton biomass dominated by picophytoplankton (*27*). Despite overall low biomass, microbial food webs in the GOM support high biodiversity of mesozooplankton and micronekton (*28*), as well as several economically important fisheries (*29*). At times, nutrient runoff from terrestrial sources promotes eutrophication, resulting in coastal hypoxic zones that are more acidic (*30*, *31*). Coastal eutrophication combined with physical upwelling of new inorganic nutrients onto the shelf can also enhance formation of harmful algal blooms (HABs), particularly along the western coast of Florida (*32*, *33*) and in other coastal regions in the southern Gulf (*34*). HABs pose a threat to marine ecosystems in the GOM and can negatively impact local economies (*35*). While OA has resulted in observable changes in ocean chemistry in the GOM (*24*), research on the impacts of OA and warming on marine microbes has not been well explored. Most microbial genomics studies have been localized to specific regions or depths (*36–39*) or focused on responses of microbes to natural disturbances, like oil spills, in the northern Gulf (*40*, *41*). This lack of spatial biological sampling has made it difficult to characterize environmental drivers of diverse microbes in the GOM (*24*), including OA parameters (e.g., pH, DIC, and *p*CO_2_), and impedes our ability to understand how microbial communities may shift in the future.

Here, we performed the first basin-scale DNA metabarcoding survey of protists, Bacteria, and Archaea in the GOM as part of the fourth Gulf of Mexico Ecosystems and Carbon Cycle (GOMECC-4) cruise that sailed from late summer to early fall of 2021. Overall, we collected 481 discrete DNA samples from 51 (out of 141) stations, encompassing 16 inshore–offshore transects and up to three depths per site that corresponded to the surface, deep chlorophyll maximum (DCM), and near bottom (Fig. 1A). Amplicon metabarcoding was performed to reveal population dynamics of protists (18S SSU rRNA gene, V9 region) and prokaryotes (16S SSU rRNA gene, V4–V5 region). We constructed generalized linear models (GLMs) for major microbial groups in the photic zone to gain insight into group-specific environmental correlates, including carbonate system parameters. These GLMs were applied to all GOMECC-4 sites, including those where DNA samples were not collected, to expand spatial distributions of microbial groups in the GOM. Finally, we performed indicator analysis based on profiles of DIC and total alkalinity (TA) in the photic zone to identify microbes that were potential indicators of more or less acidic waters (based on TA:DIC ratios). This study provides an important baseline for microbial OA research in the GOM that will guide future DNA sampling efforts in this region and contribute to our growing knowledge on the potential responses of marine microbes to climate change.

**Figure 1:**
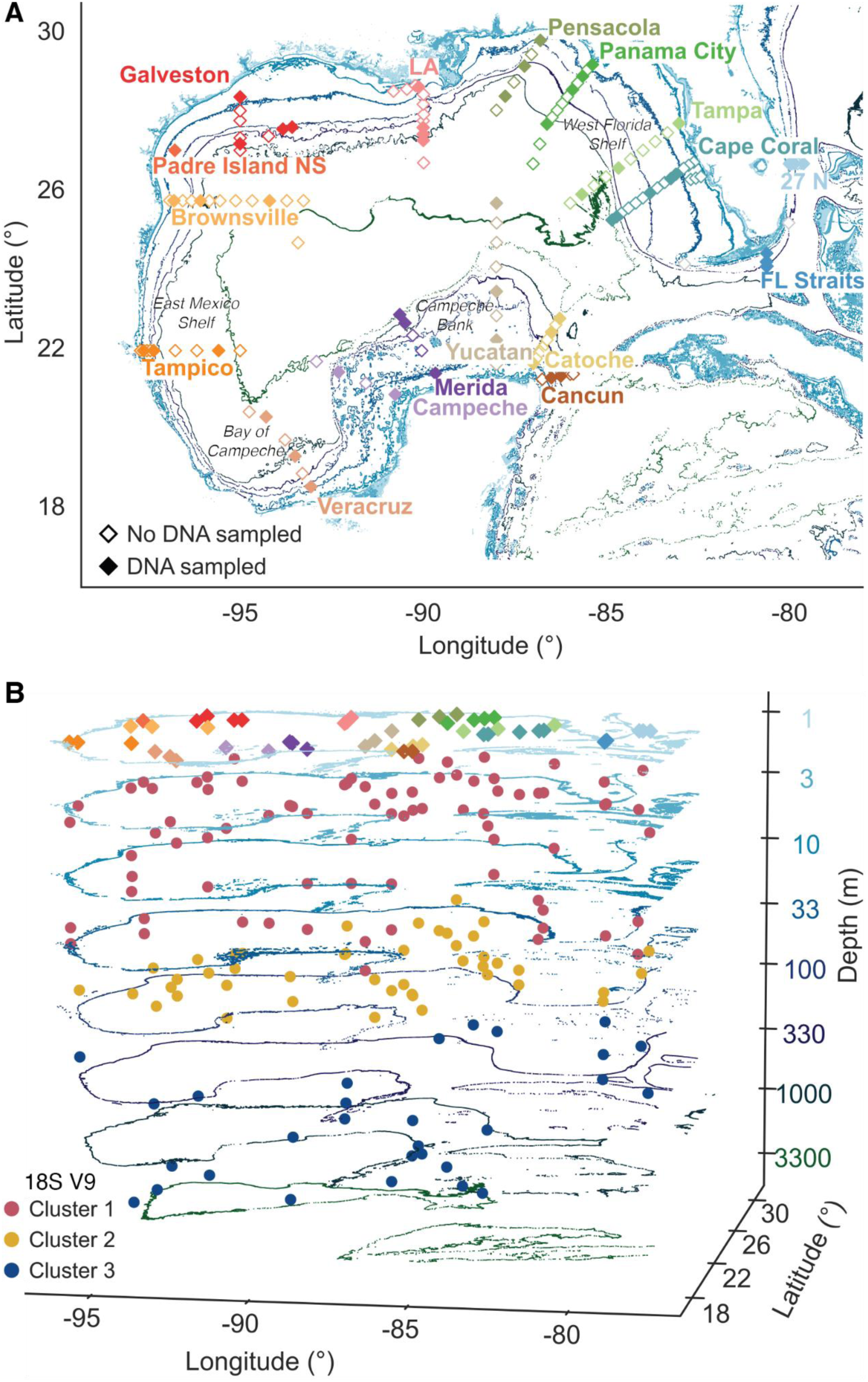
Vertical and horizontal DNA sampling across the GOM. **(A)** Map of all sites sampled on GOMECC-4. Sites are colored by transect and indicate instances where DNA was (filled) or was not (empty) sampled. Samples at all sites were collected in triplicate. Environmental metadata was collected from all stations. Contour lines indicate depth in the GOM and correspond to the right y-axis in panel **B**. Transects are also labeled to match the color of stations along a given transect. Samples were collected counterclockwise in the Gulf starting at the 27°N line. **(B)** Map displaying depth-related position (log scale) of samples across the GOM. Stations are colored by transect at the surface, matching transect colors in panel **A**. Samples with depth are colored by their clusters (Clusters 1–3) that were determined via hierarchical clustering of Aitchison distances and largely reflected depth in the water column. Cluster 1 generally corresponded to shelf waters at all depths and in the open GOM at the surface (photic zone; n = 235), Cluster 2 represented sites in the DCM in the open GOM (DCM; n = 137), and Cluster 3 was confined to open ocean sites in deep waters (aphotic zone; n = 89). Clustering was similar between 18S (shown) and 16S samples (fig. S16).

## Results and Discussion

### Microbial population dynamics in the GOM

We obtained a total of 8,312 sequences on average per sample (range: 3,322–16,483) from 18S metabarcoding, resulting in 13,632 protist amplicon sequence variants (ASVs) identified throughout all GOM samples. In comparison, we obtained an average of 12,963 sequences per sample (range: 5,056–28,620) for 16S metabarcoding which were assigned to 41,876 total prokaryotic ASVs. Though significant to community composition (*P* < 0.01), factors like transect, location on the shelf (< 200 m) vs. open ocean (> 200 m), and categorical depth had low explanatory power on their own (PERMANOVA *R* = 0.03–0.2). As depth is a well-known driver of global marine microbial communities (*42–46*), we performed hierarchical clustering of microbial composition to better control for the impact of depth on subsequent spatial analyses. This revealed separation of DNA samples into three clusters (Clusters 1–3), similar for both marker gene regions, that reflected depth of samples in the water column on the continental shelf and/or in open ocean GOM regions (Fig. 1B; fig. S1). For instance, Cluster 1 mainly consisted of samples collected on the shelf at all depths and offshore at the surface layer, with all samples located in the photic zone (2–99 m). Cluster 2 samples were mainly from the DCM (2–124 m) in more stratified open ocean regions of the GOM, while Cluster 3 samples largely represented meso-to bathypelagic waters (135–3,326 m) in the open ocean that were confined to the aphotic zone (Fig. 2A–B; Fig. 3A–B). Though Clusters 1–2 were both in the photic zone (upper ∼150 m), and had some overlap (Fig. 2B), they were separated into distinct clusters based on their composition that reflected total depth in the water column and shifts in physicochemical variables (fig. S2). In our case, clustering of DNA samples allowed us to better explore microbes, and their relationships with environmental variables, within distinct spatial habitats they occupy in the GOM.

**Figure 2:**
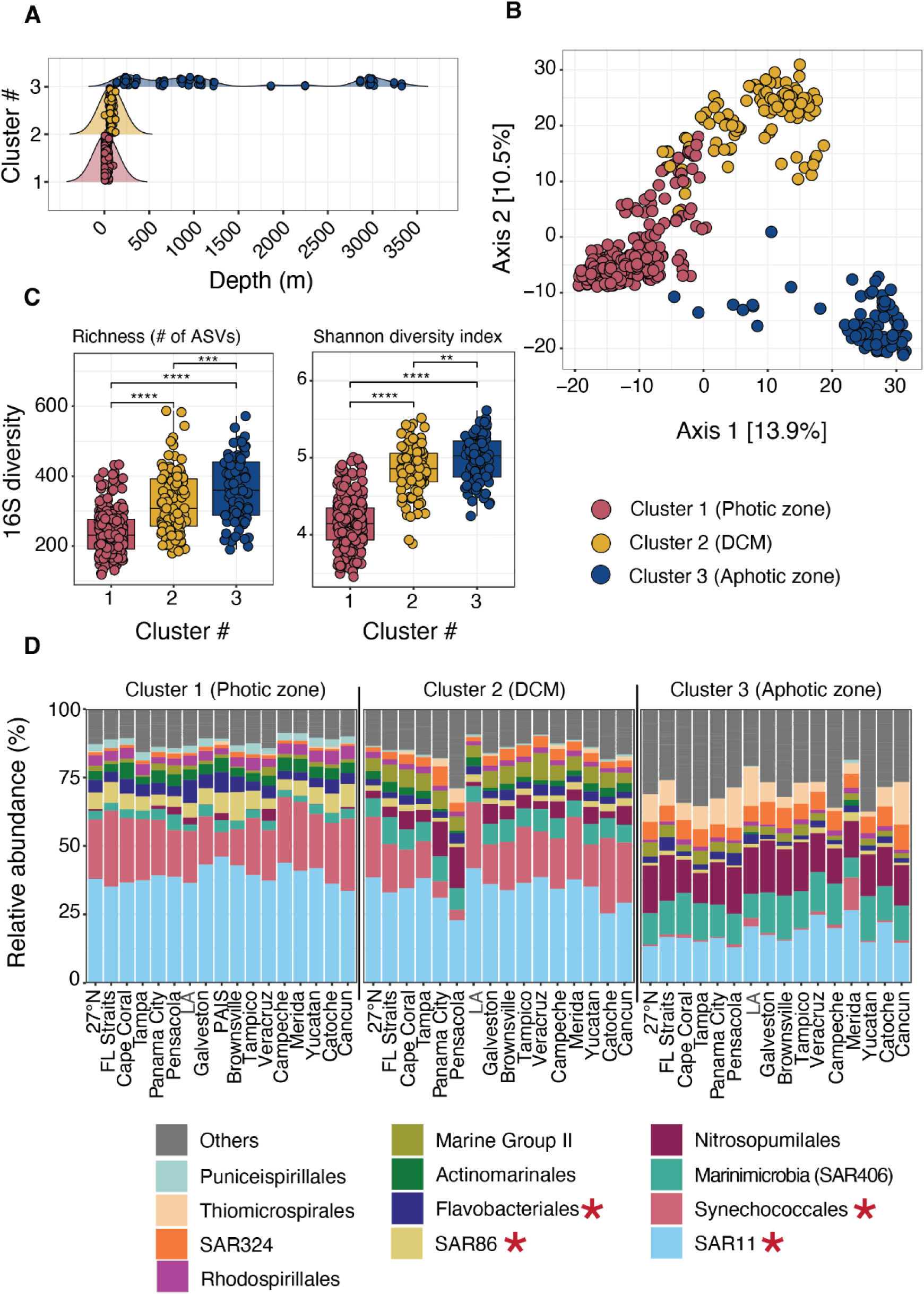
Bacterial and archaeal community dynamics in the GOM from 16S metabarcoding. **(A)** Ridgeline plots showing the depth distribution of samples within each cluster (Clusters 1–3). Clusters were determined via hierarchical clustering of Aitchison distances: Cluster 1 (photic zone), Cluster 2 (DCM), and Cluster 3 (aphotic zone). **(B)** Principal coordinates analysis of Aitchison distances, with samples colored by their respective clusters. **(C)** Mean observed richness (# of ASVs) and Shannon diversity index for Clusters 1–3, with points representing individual samples. Significant differences between clusters were determined with Wilcoxon tests (** *P* < 0.01, *** *P* < 0.001, **** *P* < 0.0001). **(D)** Stacked bar plots of mean relative abundance (%) at the order level in each sampling transect and faceted by cluster. Transects are ordered on the x-axis based on the order of sampling (counterclockwise) on GOMECC-4, except for FL straits and Cape Coral that were sampled last but grouped spatially with other FL lines. Bar plots display the top 12 most relatively abundant groups over all samples (“others” in gray). Taxonomy was assigned via the SILVA database. Generalized linear models focused on the top four most relatively abundant groups in Cluster 1 (red asterisks). Models for Synechococcales were constructed at the genus level to discriminate between *Prochlorococcus* and *Synechococcus*. LA = Louisiana and PAIS = Padre Island National Seashore. Transects have the same labels in all subsequent plots.

**Figure 3:**
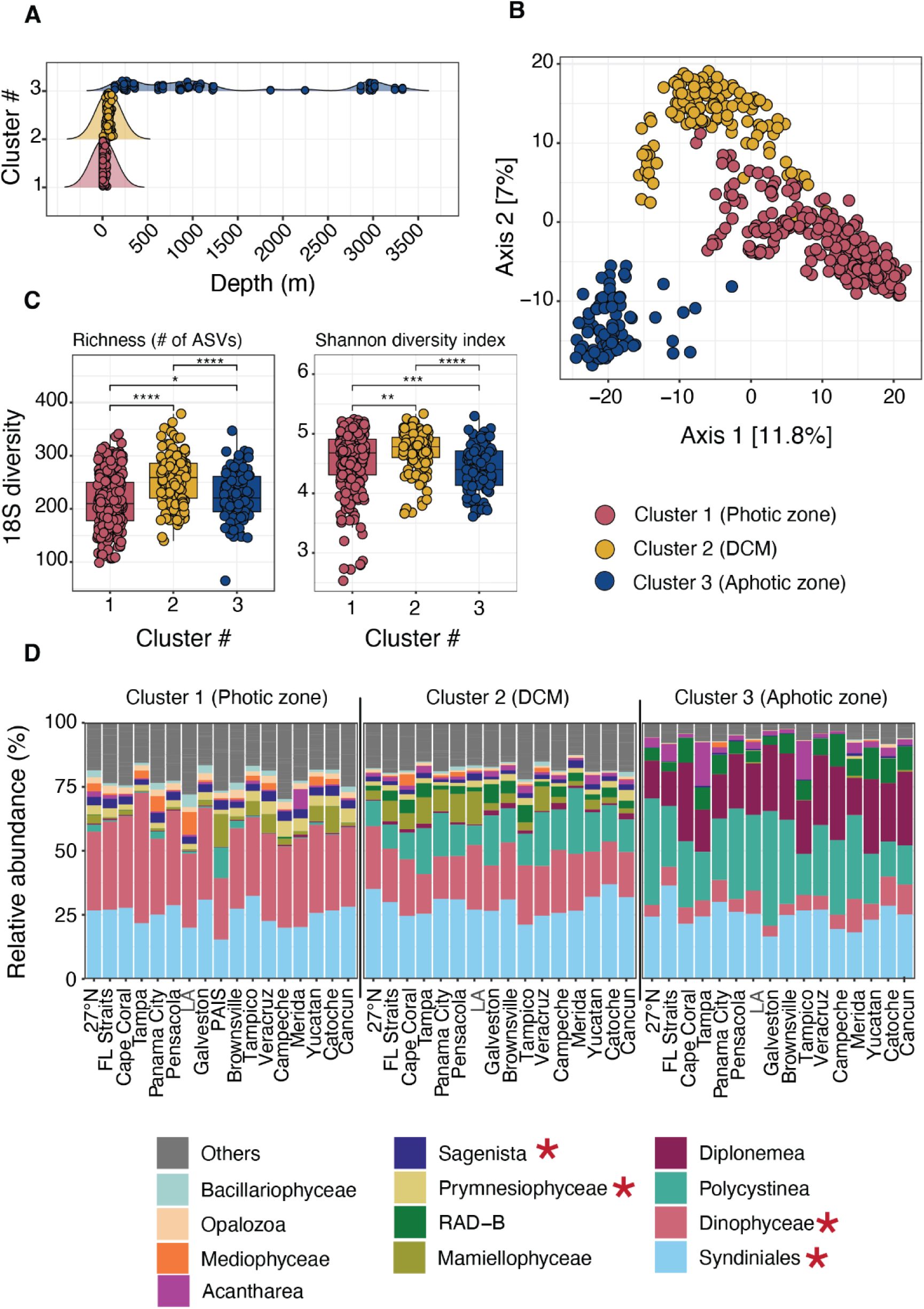
Protist community dynamics in the GOM from 18S metabarcoding. **(A)** Ridgeline plots showing the depth distribution of samples within each cluster (Clusters 1–3). Clusters reflected depth along the shelf and open GOM and closely resembled clustering of 16S samples**. (B)** Principal coordinates analysis of Aitchison distances, with 18S samples colored by cluster. **(C)** Mean observed richness (# of ASVs) and Shannon diversity index for Clusters 1–3, with points representing individual samples. Significant differences between clusters were determined with Wilcoxon tests (* *P* < 0.05, ** *P* < 0.01, *** *P* < 0.001, **** *P* < 0.0001). **(D)** Stacked bar plots of mean relative abundance (%) at the class level in each sampling transect and faceted by cluster. Transects are ordered the same as in Fig. 2. Bar plots display the top 12 most relatively abundant groups over all samples (“others” in gray). Protist taxonomy was assigned via the PR2 database. Generalized linear models focused on the top four most relatively abundant groups in Cluster 1 (red asterisks).

Microbial communities in the GOM were more species-rich and diverse in the DCM and aphotic zone (Fig. 2C; Fig. 3C), consistent with vertical profiles from other oceanic regions (*43*, *46*, *47*). Higher richness and diversity with depth may be the result of microbes utilizing a broad spectrum of sinking organic matter, exerting alternative metabolic strategies (redox reactions), and/or forming diverse trophic relationships with other organisms to exploit such habitats (*48*). Alpha diversity was stable along sampling transects in the photic zone for 16S (fig. S3) and 18S samples (fig. S4), with higher variability in the DCM and aphotic zone. For example, microbial diversity in the aphotic zone steadily decreased from coastal Florida (27°N line) to regions near the Mississippi River outflow, increasing thereafter from Brownsville to Cancun (fig. S3).

Shifts in taxonomy between clusters were in line with depth-related microbial dynamics seen previously in the GOM (*37*, *38*, *49*) and on a global scale (*42*, *43*, *48*). Among prokaryotes, the photic zone and DCM were dominated by common heterotrophic bacteria, such as SAR11, SAR86, and Flavobacteriales (Fig. 2D; fig. S5). Autotrophic cyanobacteria within the order Synechococcales also had high relative abundance in the photic zone (Fig. 2D; fig. S5), particularly *Prochlorococcus* and *Synechococcus* (fig. S6), both genera known to dominate primary production in the GOM (*27*). Prokaryotic communities shifted dramatically in the aphotic zone, with higher relative abundance of metabolically diverse taxa that are endemic to deeper waters (*48–50*), including nitrous oxide-reducing Marinimicrobia (SAR406), ammonia-oxidizing Nitrosopumilales, and sulfur-oxidizing Thiomicrospirales (Fig. 2D; fig. S5). These microbes use redox reactions to acquire energy in less oxygenated waters (*48*), such as those found in the mesopelagic zone (∼200–800 m) in the GOM (fig. S2), and likely contributed to increased richness of prokaryotic communities observed with depth (Fig. 2C). Certain 16S groups varied at more resolved taxonomic levels between clusters. For example, *Prochlorococcus* became more relatively abundant in the DCM, while SAR11 clade II increased in the aphotic zone relative to other SAR11 clades (fig. S7). Similar patterns have been observed elsewhere for *Prochlorococcus* (*51*) and SAR11 (*48*, *52*), and reflect potential environmental niche partitioning through the water column. High abundance of SAR11 clade II in the mesopelagic has recently been observed in the Pacific Ocean (*48*), which may indicate particle association among SAR11 that may be more common than previously thought.

Protist biodiversity was dominated by Dinophyceae, Syndiniales, Prymnesiophyceae, and Sagenista in the photic zone and DCM, transitioning to Radiolaria (Polycystinea and RAD-B) and Diplonemea in the aphotic zone (Fig. 3D). Dinophyceae and Prymnesiophyceae are common in pelagic waters, including in the GOM (*36*, *53*), and occupy important functional roles as grazers (and mixotrophs) in microbial food webs (fig. S5). Sagenista was also abundant in the photic zone (Fig. 3D), a group of common, yet still uncultured heterotrophic protists that have important ecological roles (*54*). Other class level protist groups that were common in the GOM in summer–fall, like Mamiellophyceae (Chlorophyta) and Mediophyceae (Stramenopiles), varied more greatly across transects in the photic zone and DCM (Fig. 3D). Radiolarians dominated relative abundance in mesopelagic samples (Fig. 3D). While these organisms remain largely uncultivated and hard to study, they are key members of deep ocean food webs, forming endosymbiotic relationships with other microorganisms (fig. S5) and contributing to the export of carbon and biogenic silica (*55*).

DNA metabarcoding also reinforced the importance of obligate parasites within the group Syndiniales at all depths in the water column (*42*, *56*, *57*), including at the basin scale in the GOM (Fig. 3D; fig. S5). The prevalence of Syndiniales may be attributed to their wide host range, active (and passive) export on sinking particles, and depth related niche partitioning (*56*, *58*). We observed vertical shifts within Syndiniales at the clade level in our samples that aligned with prior observations (*45*, *56*). For instance, there was a shift from Syndiniales Group-I Clades 1 and 4 in the photic zone to other clades, like Group-II Clade 7 and Group-I Clade 2 in the aphotic zone (fig. S8). Radiolaria also varied between clusters, with certain members of Polycystinea (e.g., *Heliosphaera* and *Pterocorys*) increasing in relative abundance from the DCM to the aphotic zone (fig. S8). Diplonemea were also dominant in the GOM aphotic zone (Fig. 3D). Though enigmatic, Diplonemea have been found globally in mesopelagic waters (*57*) and likely represent important consumers of picoplankton and bacteria in these environments (*59*).

### Generalized linear models reveal group-specific environmental correlates

We used generalized linear models (GLMs) with either Poisson or negative-binomial error distributions to identify potential explanatory variables of major 16S and 18S taxonomic groups in the GOM. GLMs account for multiple predictor variables (factors) and have been applied to ecological count (and proportional) data of higher trophic level marine organisms (*60*, *61*). Here, we applied GLMs to microbial metabarcoding data, allowing us to observe predictor variables and their relation to group-specific relative abundance measured spatially in the photic zone (Fig. 2D; Fig. 3D). We focused our models on the photic zone (Cluster 1), primarily because most factors were collinear in the DCM and aphotic zone (table S1). Collinearity among variables can result in models being less statistically reliable and confound model interpretation (*62*). Eight of fifteen environmental variables were initially selected for models and included temperature, salinity, dissolved oxygen (O_2_), nitrate (NO_3_), ammonium (NH_4_), phosphate (PO_4_), dissolved inorganic carbon (DIC), and total pH recalculated to in situ temperatures (Table 1). Many parameters related to OA that were measured or derived (e.g., total alkalinity, *p*CO_2_, carbonate ion concentration, and aragonite saturation) were strongly collinear to each other and temperature (Spearman *r_s_* > 0.7 or < –0.7), and thus were excluded from initial models (table S1). Environmental conditions in the GOM surface were typical for this time of year (*63*, *64*). For instance, offshore waters were warm (> 28 °C) and nutrient-limited (e.g., NO_3_ < 0.1 µmol kg^−1^), while coastal regions had higher nutrient concentrations, including near the Mississippi River outflow (Table 1; fig. S2). DIC was highest in the southern GOM and onto the Campeche Bank (> 2050 µmol kg^−1^), while pH often increased from the shelf to open ocean regions of the Gulf (Table 1; fig. S2).

**Table 1:**
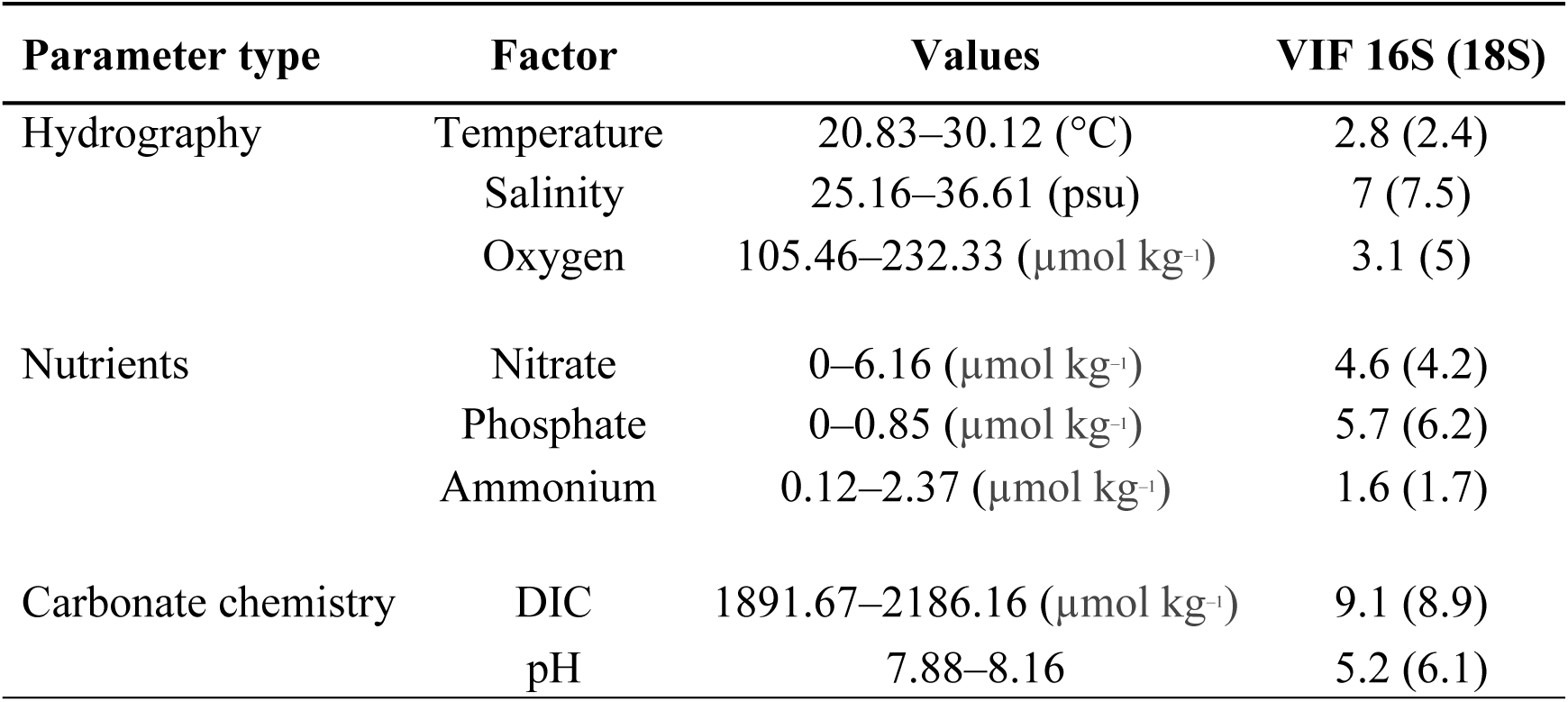
Environmental factors used in microbial models. Factors were grouped into parameter type and chosen for initial GLMs based on Spearman correlations (table S1) and low variance inflation factors (VIF < 10) to mitigate collinearity among predictor variables. VIFs varied slightly between 16S and 18S (in parentheses) due to differences in sample size (n = 274 for 16S; n = 235 for 18S) following clustering analysis. Triplicate samples were included in models. Datasets clustered similarly, as evidenced by a similar range in the predictor values. Initial factors were used to construct group-specific models.

| Parameter type | Factor | Values | VIF 16S (18S) |
| --- | --- | --- | --- |
| Hydrography | Temperature | 20.83–30.12 (°C) | 2.8 (2.4) |
|  | Salinity | 25.16–36.61 (psu) | 7 (7.5) |
|  | Oxygen | 105.46–232.33 (μmol kg<sup>-1</sup>) | 3.1 (5) |
| Nutrients | Nitrate | 0–6.16 (μmol kg<sup>-1</sup>) | 4.6 (4.2) |
|  | Phosphate | 0–0.85 (μmol kg<sup>-1</sup>) | 5.7 (6.2) |
|  | Ammonium | 0.12–2.37 (μmol kg<sup>-1</sup>) | 1.6 (1.7) |
| Carbonate chemistry | DIC | 1891.67–2186.16 (μmol kg<sup>-1</sup>) | 9.1 (8.9) |
|  | pH | 7.88–8.16 | 5.2 (6.1) |

Microbial groups differed in the type and number of variables that significantly contributed to the final models (Table 2). Pseudo *R*^2^ values produced from GLMs ranged from 0.26–0.80 (Table 2), though several other methods confirmed appropriate model fit. First, model simulations fit the data well (Fig. 4C; Fig. 5C) and the standardized residuals were normally distributed for all groups (Kolmogorov–Smirnov, *P* > 0.05), except for Flavobacteriales. This was further supported by significant and often strong positive correlations (Pearson *R* = 0.31– 0.90; *P* < 0.01) between test and model-trained relative abundance data for all groups (fig. S9; fig. S10), with example plots shown for SAR11 (Fig. 4D) and Syndiniales (Fig. 5D). Explanatory variables like temperature, DIC, and pH had individual, and often significant (*P* < 0.05) effects on relative abundance that varied among major 16S (Fig. 4A–B) and 18S groups (Fig. 5A–B). Through this approach, we examined individual model terms, focusing primarily on those related to ocean change, and explored their relationships with group-specific relative abundances in the GOM.

**Figure 4:**
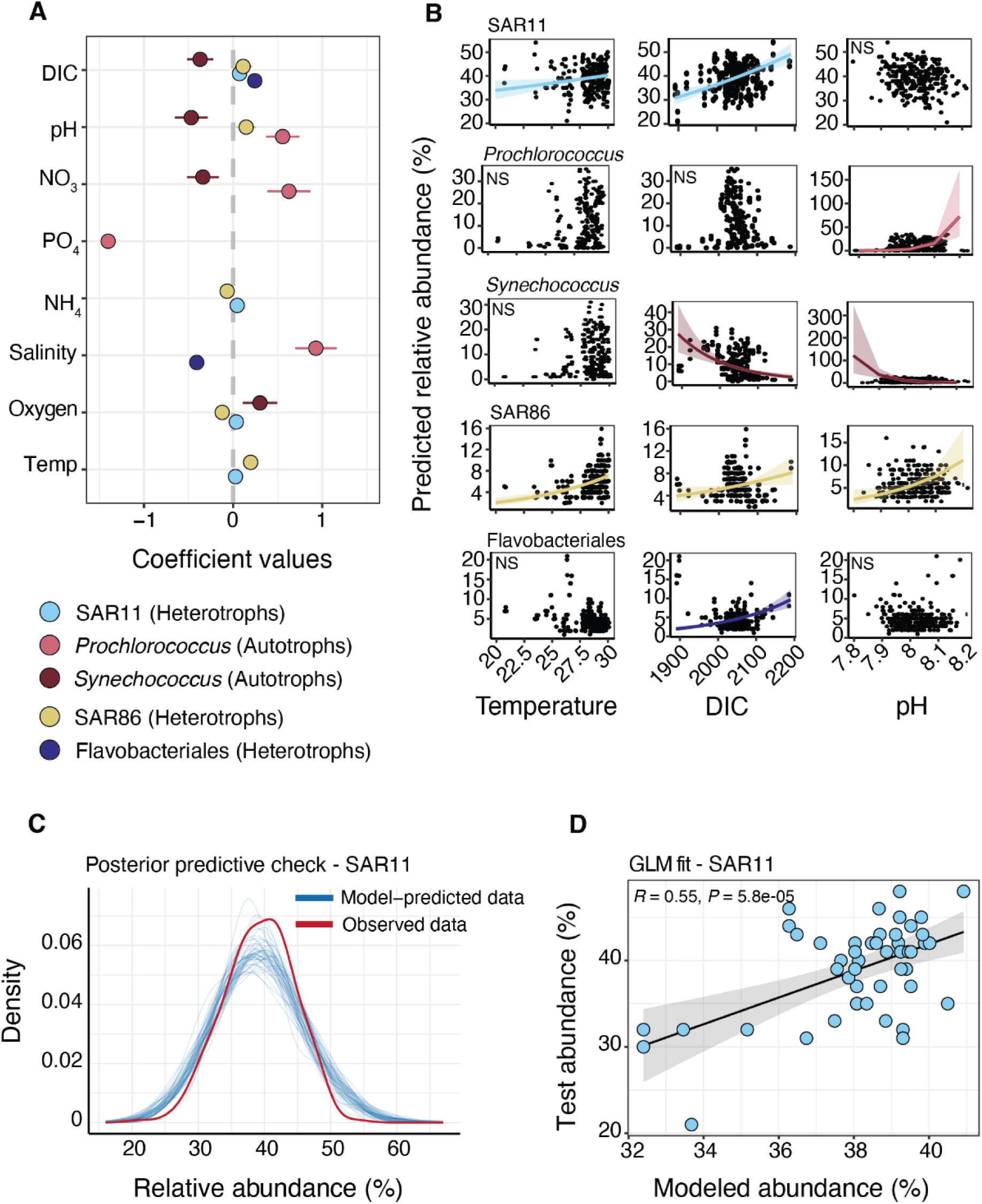
Generalized linear models of major 16S taxa reveal group-specific environmental drivers in the photic zone. **(A)** Scaled model coefficients (± 2 standard deviations) of predictor variables (environmental factors) that were significant to the final model (based on AIC values). Models were constructed with group-specific relative abundance as the response variable. The most relatively abundant 16S groups were modeled, which included heterotrophs and autotrophs. Models were generated at the order level, except for cyanobacteria (Synechococcales), where separate models were run for *Prochlorococcus* and *Synechococcus*. Only covariates that were statistically significant to a given model were plotted. **(B)** Predicted response estimates (relative abundance) and 95% confidence intervals (CIs) of major 16S groups to temperature, DIC, and in situ pH. NS = not significant. **(C)** An example of a posterior predictive plot, highlighting the fit of observed vs. model-predicted relative abundance for the final SAR11 model. The model-predicted data was simulated with 50 bootstraps and followed a similar trend as the observed data. **(D)** Pearson correlation (with 95% CI) between SAR11 test and modeled relative abundance to estimate model fit. Predicted abundance was derived from the final SAR11 model using a subset of the data (80%; 219 samples) and correlated to test data that was left out (20%; 48 samples). Model fit of other major 16S groups is shown in fig. S9.

**Figure 5:**
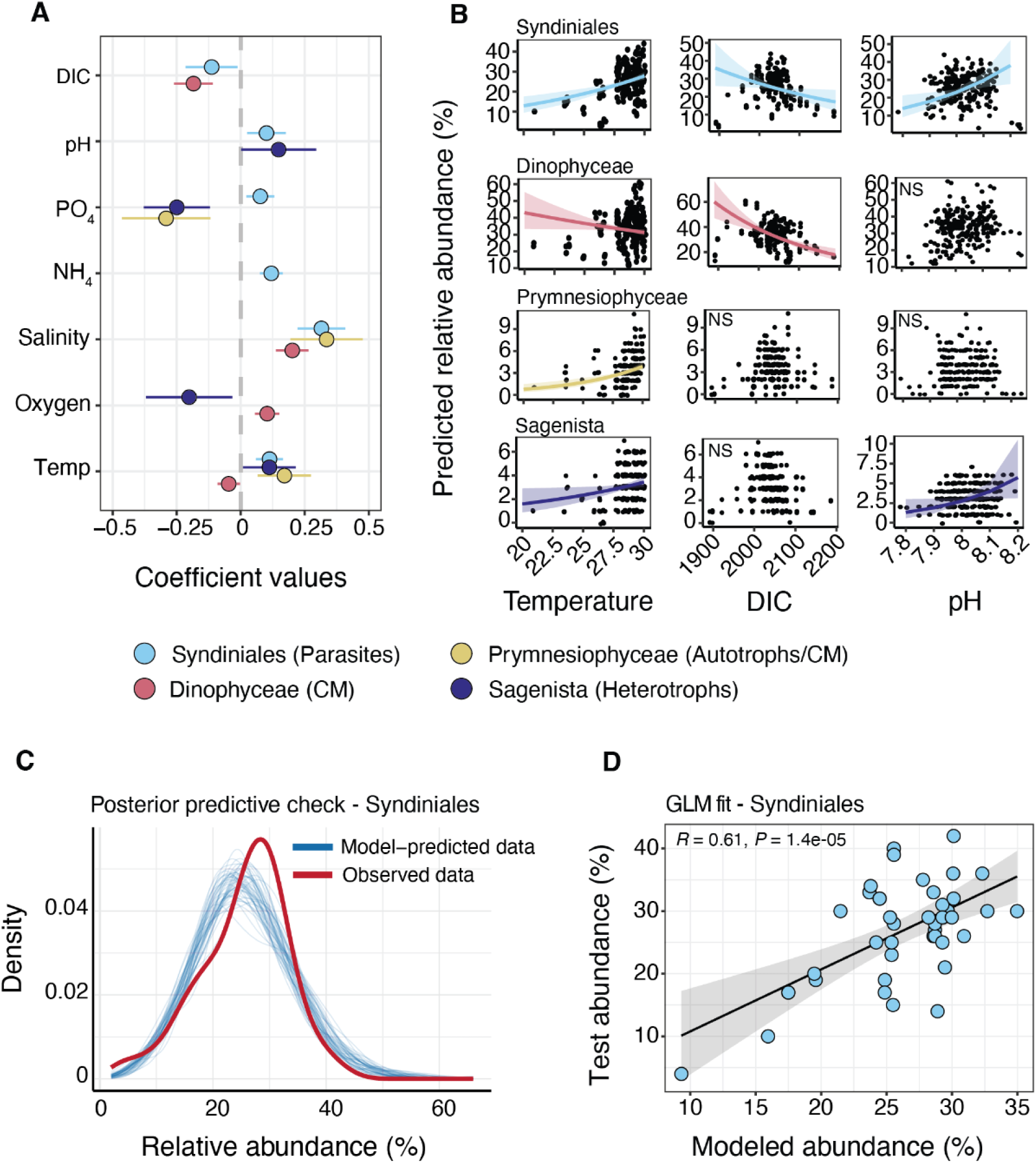
GLMs of major 18S taxa reveal group-specific drivers in the photic zone. **(A)** Scaled model coefficients (± 2 standard deviations) of predictor variables (environmental factors) that were significant to the final model (based on AIC values). Models were constructed with group-specific relative abundance as the response variable. The top four most relatively abundant 18S groups were modeled separately, spanning constitutive mixotrophs (CM), parasites, autotrophs, and heterotrophs. Covariates that were not statistically significant to a given model are not shown. **(B)** Predicted response estimates (relative abundance) and 95% confidence intervals (CIs) of major 18S groups to temperature, DIC, and pH. NS = not significant. **(C)** An example of a posterior predictive plot, highlighting the fit of observed vs. model-predicted relative abundance for the final Syndiniales model. The model-predicted data was simulated with 50 bootstraps and followed a similar trend as the observed data. **(D)** Pearson correlation (with 95% CI) between Syndiniales test and modeled relative abundance. Predicted abundance was derived from the final Syndiniales model using a subset of the data (80%; 187 samples) and correlated to test values that were left out (20%; 43 samples). Model fit of other major 18S groups is shown in fig. S10.

**Table 2:** Final models for major microbial groups in the photic zone. Protists were examined at the class level and prokaryotes at the order level. GLMs were constructed for *Prochlorococcus* and *Synechococcus*. Models were run either with negative binomial (neg bin) or Poisson distributions. Variables that were significant to the final model (*P* < 0.05) are shown for each group and reflect stepwise selection based on Akaike Information Criterion (AIC). See Table 1 for the full list of variables considered. Pseudo *R*^2^ values are shown as a proxy for model fit, though standardized residuals and validation tests confirmed model fit. Temp = temperature (°C); Sal = salinity; Oxy = oxygen (µmol kg^−1^); PO_4_ = phosphate (µmol kg^−1^); NO_3_ = nitrate (µmol kg^−1^); NH_4_ = ammonium (µmol kg^−1^); DIC = dissolved inorganic carbon (µmol kg^−1^).

| Group | Taxonomy | GLM | Type | $R^2$ |
| --- | --- | --- | --- | --- |
| Protists | Dinophyceae | Temp + Sal + Oxy + DIC | Neg Bin | 0.5 |
| | Syndiniales | Temp + Sal + $\text{PO}_4$ + $\text{NH}_4$ + pH + DIC | Neg Bin | 0.57 |
| | Sagenista | Temp + Oxy + $\text{PO}_4$ + pH | Poisson | 0.28 |
| | Prymnesiophyceae | Temp + Sal + $\text{PO}_4$ | Poisson | 0.6 |
| Prokaryotes | SAR11 | Temp + Oxy + $\text{NH}_4$ + DIC | Poisson | 0.26 |
| | *Synechococcus* | Oxy + $\text{NO}_3$ + pH + DIC | Neg Bin | 0.38 |
| | *Prochlorococcus* | Sal + $\text{PO}_4$ + $\text{NO}_3$ + pH | Neg Bin | 0.8 |
| | SAR86 | Temp + Oxy + $\text{NH}_4$ + pH + DIC | Poisson | 0.33 |
|  | Flavobacteriales | Sal + DIC | Poisson | 0.64 |

Our model findings often supported prior physiological responses for certain microbial groups that have been revealed in field and culture experiments. While changes in relative abundance data with any given environmental factor does not necessarily translate to physiology, applying DNA metabarcoding to OA research can help to verify existing trends and produce new hypotheses for future testing on a wide range of microorganisms (*22*, *23*). We found that temperature had a positive effect on the relative abundance of SAR11 and SAR86 in our models (Fig. 4A–B). Experimental evidence suggests that warmer conditions may favor increased biomass of small, oligotrophic bacteria, like SAR11 and SAR86, that have low nucleic acid content (*65*). In general, warming is thought to promote increased bacterial production, biomass, and respiration, while also lowering growth gross efficiency (*20*). DIC had a positive effect on SAR11, SAR86, and Flavobacteriales in group models (Fig. 4A–B), which together with temperature effects, may imply a favorable response among these taxa to continued OA and warming in this region. It is important to note that heterotrophic bacteria will also be influenced by indirect changes in plankton composition, dissolved organic matter (DOM) availability and quality, and trophic interactions (*21*, *66*). These factors may outweigh direct OA effects in natural communities and will be important to incorporate into future climate model predictions of bacterial diversity and composition.

We observed contrasting effects of pH on the relative abundance of *Prochlorococcus* vs. *Synechococcus* in the photic zone (Fig. 4A–B). For example, in situ pH had a strong and positive effect on *Prochlorococcus*, implying a negative response to lower pH (more acidic) conditions. An opposite trend was observed for pH in the *Synechococcus* model (Fig. 4A–B); however, DIC also had a negative effect on *Synechococcus*, confounding model inference. Fu et al. (2007) noted that combined effects of high *p*CO_2_ and temperature significantly increased growth rates, photosynthetic capacity, and cellular pigment levels of *Synechococcus* but not *Prochlorococcus*. Mesocosm work in the subtropical North Atlantic also indicated a positive response of *Synechococcus* to high *p*CO_2_ (*67*), though others have noted small or insignificant physiological shifts to changing conditions (*68*). On a global scale, ecological niche models predict increased *Prochlorococcus* and *Synechococcus* biomass to ocean warming (via flow cytometry), particularly in low latitude regions like the GOM, where these taxa already dominate plankton biomass (*69*). Such niche models have not considered pH (or DIC), which we show may be significant predictors. Though not resolved at the ecotype level in our amplicon dataset, individual cyanobacterial strains or ecotypes will likely respond differently to future conditions, as well as be influenced by indirect changes in top-down pressure (grazing or viral lysis), nutrients, or sunlight (*70*, *71*). Multiple ecotypes have already been discovered for *Prochlorococcus* in the ocean, with evidence of different nutrient uptake rates, light preferences, and thermal optima that shape population dynamics (*51*, *72*). Additional field and laboratory work is needed to identify responses among microbes at the species or ecotype level to support accurate model predictions (*13*) and reveal underlying physiological mechanisms.

Future OA and warming is predicted to favor small phytoplankton, like picoeukaryotes, that can more efficiently exploit oligotrophic and nutrient-limited waters (*11*, *12*, *14*), primarily due to their larger cell surface to volume ratios that promote resource acquisition. Though for many protists, the effects of OA and warming are less clear. This is especially true for Syndiniales and Sagenista, enigmatic protist parasites and grazers that have seldom been considered with respect to climate change. In our models, temperature had a significant and positive effect on the relative abundance of Syndiniales and Sagenista (Fig. 5A–B). Temperature is often thought to enhance physiological rates (*73*), which may include microzooplankton grazing and parasitism; however, temperature relationships are hard to predict and can often be confounded by other factors, like host or prey composition, that can dictate mortality rates. We found that pH had a positive effect on Syndiniales and Sagenista, while DIC had a negative impact on Syndiniales relative abundance (Fig. 5A–B). This implied a negative response among these groups to more acidic conditions in the GOM at this time. Therefore, models that include only temperature or pH may result in different outcomes for certain plankton groups (*74*), potentially misleading how we interpret (and predict) their responses to climate change.

Dinophyceae were also prevalent in the photic zone on GOMECC-4 (Fig. 3D). It is well understood that dinoflagellates are central to the microbial loop in oligotrophic regions, often exhibiting mixotrophy and representing a key link between primary production and higher trophic levels (*29*, *53*). We found that temperature and DIC had significant and negative effects on Dinophyceae relative abundance (Fig. 5A–B), implying a negative response to warmer and/or more acidic conditions in the GOM. Similar findings on dinoflagellates have been observed in a mesocosm study (*67*), though others have found dinoflagellates to benefit from or be less sensitive to warming or increased *p*CO_2_ concentrations (*23*, *71*, *74*, *75*). Dinoflagellates often exhibit mixotrophy, and so favorable responses to OA among this group may be indicative of increased consumption of common prey (picoeukaryotes and cyanobacteria) that tend to grow faster under such conditions (*75*). As is the case with many protists, dinoflagellates are extremely diverse, not only phylogenetically but also in terms of their size, physiology, and trophic modes (*76*). Therefore, it will likely be challenging to define a unified response for Dinophyceae to changing ocean conditions. Future work that merges DNA metabarcoding with more targeted approaches, like single-cell genomics or qPCR, will help to shed light on species sensitivity, interactions, and drivers that would otherwise be overlooked.

In addition to temperature and carbonate chemistry parameters, other factors like nutrients, salinity, and oxygen had significant effects on prominent 16S and 18S groups in the GOM (Fig. 4A; Fig. 5A). As an example, limiting nutrients like NO_3_ and PO_4_ had negative effects on the relative abundance of *Synechococcus* and *Prochlorococcus*, respectively (Fig. 4A). This may be related to the ability of cyanobacteria to uptake nutrients at low concentrations in surface waters (*72*). *Synechococcus* are thought to exploit low NO_3_ concentrations in the GOM by maintaining a shallower distribution in the water column (*64*), relying on regenerated sources of NO_3_ via nitrification (*29*). Salinity was also an important variable in our models, with positive effects on the relative abundance of *Prochlorococcus*, Syndiniales, Dinophyceae, and Prymnesiophyceae, as well as a negative effect on Flavobacteriales (Fig. 4A; Fig. 5A). Salinity is a known driver of bacterial and plankton distribution and diversity in the GOM (*37*, *39*, *77*). This is particularly evident in the northern GOM, where plankton biomass and composition are often driven by salinity-induced stratification (and nutrient availability) that result from riverine discharge via the Mississippi–Atchafalaya system, as well as by climatic processes, like the El Niño–Southern Oscillation (*77*, *78*). Here, interpreting the role of salinity or nutrients in driving specific microbial groups was difficult, mainly because our sampling strategy and analysis focused on large-scale spatial patterns in microbial communities that did not allow us to explore regional trends (e.g., in the northern GOM). Even so, our results emphasize the importance of including such variables to resolve microbial composition and distribution at the basin scale in the GOM.

There are several caveats to consider with our model analysis. Models constructed from amplicon data on GOMECC-4 reflected only a specific time of the year (summer–fall) and did not integrate seasonal sampling. Temperature and carbonate chemistry parameters vary seasonally in the GOM (*79*), as does the intensity and position of the Loop Current (and eddies) and nutrient input from coastal runoff, all of which will impact microbial communities (*80*, *81*). Consistent temporal sampling will be essential to better resolve microbes and their drivers over seasonal and interannual time scales (*24*). Such sustained sampling will also allow for more accurate predictions of microbial dynamics that integrate new OA data beyond the limits of GOMECC-4 measurements. We also considered GLMs for major taxonomic groups that were present in our samples (i.e., highest relative abundance), mainly to avoid issues with zero-inflation and overdispersion in the models. As a result, several groups thought to be sensitive to ocean change, like diatoms and diazotrophic cyanobacteria (*8*), were not considered here due to lower relative abundance at the basin scale. Similarly, this constrained our ability to predict model effects below the order to class level, with the exception being highly abundant cyanobacterial genera. We examined linear trends with GLMs as a simple and conservative approach to model relative abundance in the GOM. Future work may consider applying generalized additive models (GAMs) that allow for nonlinear dynamics (*82*), especially as more amplicon data is collected.

Though we tested for model fit, it is important to note that amplicon data is compositional, with relative abundance of any single group being dependent on the proportion of others (*83*). Comparing relative abundance among eukaryotic groups is also tenuous, as 18S rRNA gene copy numbers can vary greatly (2–166 copies per cell) among protists (*84*). This is especially true for alveolates (Syndiniales and Dinophyceae) and can lead to overestimation of read counts and relative abundance (*42*). However, such concerns would not necessarily discount our modeling approach that focused on groups separately and explored their relation to environmental factors. Lastly, models did not account for trophic interactions (e.g., changes in prey or host) that may vary along with changing conditions (*19*) or potential evolutionary adaptations among organisms (*85*). Nevertheless, applying GLMs to amplicon data in this study offered a first step to define multiple environmental drivers of diverse marine microbes, many of which are not easily discerned with traditional observational methods like microscopy or cytometry. Our findings are also timely for marine regions like the GOM that have lacked basin-scale sampling.

### GLMs expand microbial distributions in GOM surface waters

Final models were used to predict the relative abundance of major 16S and 18S groups at 135 surface sites on GOMECC-4 (Fig. 1A), including 84 sites where DNA was not collected. This allowed us to increase the spatial resolution of microbial sampling in the GOM at this time of year. Groups like SAR11 (Fig. 6A) and Syndiniales (Fig. 6F) were well distributed throughout the GOM, with highest relative abundance predicted offshore of Brownsville (Texas), in the Bay of Campeche, and regions on the Campeche Bank. Cyanobacteria genera were largely partitioned in the GOM based on their expected ecological niches (*86*). *Prochlorococcus* was most relatively abundant offshore in stratified and nutrient-limited waters (Fig. 6B), particularly in parts of the southern GOM. *Synechococcus* was present throughout the GOM at the surface, but relative abundance was often highest in nutrient-rich coastal regions and in a localized area in the central Gulf (Fig. 6C). Other groups like SAR86 (Fig. 6D), Prymnesiophyceae (Fig. 6H), and Sagenista (Fig. 6I) were most relatively abundant offshore in the southern GOM and onto the East Mexico Shelf, likely driven by higher temperature and DIC concentrations in these areas (fig. S2). This was supported in the model output for these taxa, where temperature and/or DIC had positive effects on relative abundance (Fig. 4A; Fig. 5A). Flavobacteriales was highest near the Mississippi River outflow (Fig. 6E), in line with strong negative effects of salinity in the model output for this group (Fig. 4A). The Mississippi River is the dominant source of freshwater into the GOM, providing nutrients and organic matter into the system (fig. S2) that can stimulate phytoplankton blooms (*81*). Though not widespread in the GOM, diatoms were most relatively abundant in the photic zone near the Mississippi River (Fig. 3D), which may have contributed to higher relative abundance of copiotrophs like Flavobacteriales that often associate with blooms and can rapidly consume DOM (*87*).

**Figure 6:**
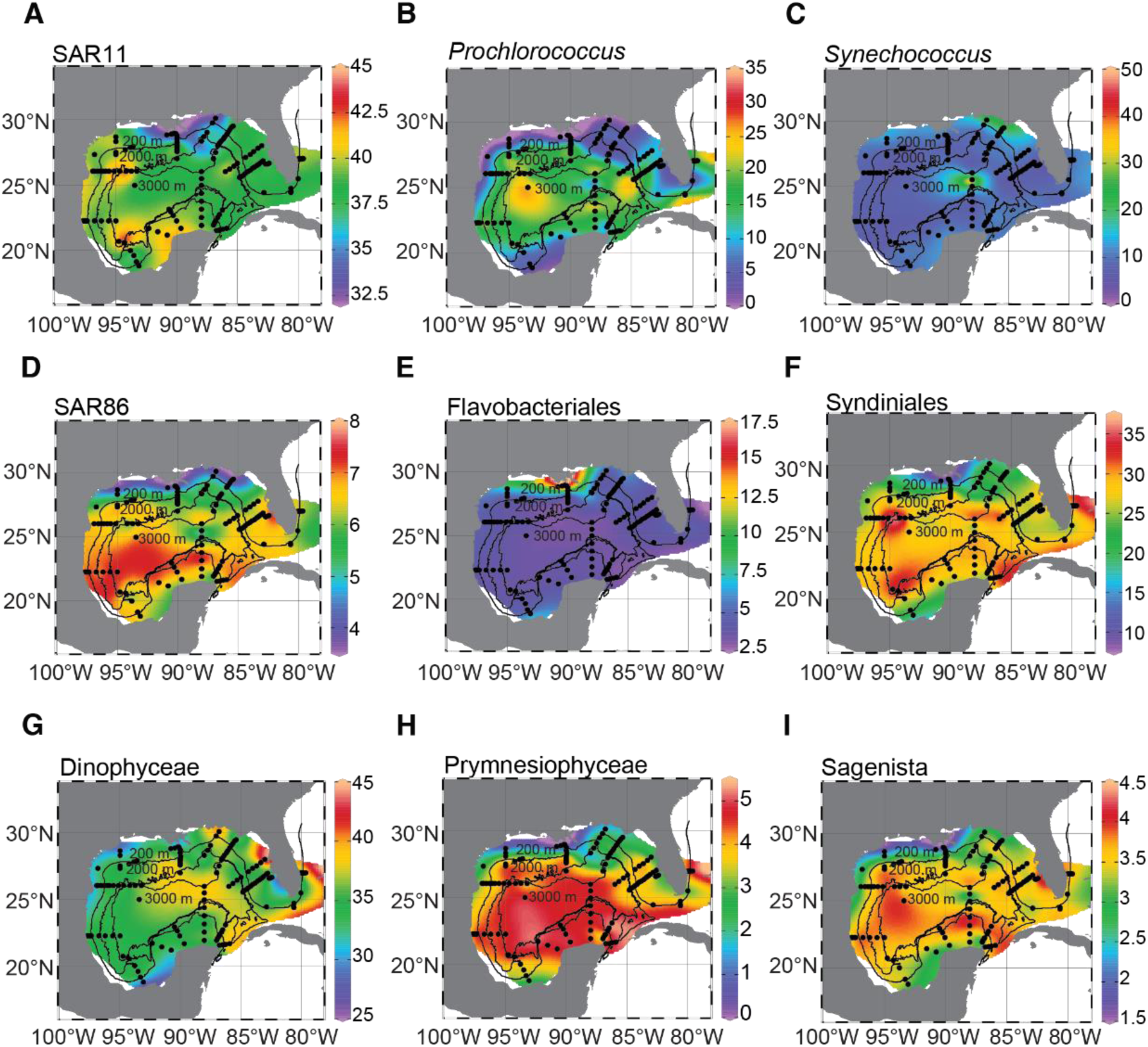
Expanding current microbial distributions in the GOM. Predicted relative abundance (%) of major 16S **(A**–**E)** and 18S groups **(F**–**I)** at 135 GOMECC-4 sites modeled with each respective GLM (from Table 2). Model results have been interpolated using DIVA interpolation in Ocean Data View. Isobaths are shown for 200 m, 2,000 m, and 3,000 m. Scales for predicted relative abundance vary by taxonomic group (on the right of each panel) but display low to high relative abundance.

Current predictions also revealed insights into the biogeography of a HAB species in the GOM. Though prevalent through most of the GOM, including in the open ocean, relative abundance of Dinophyceae was predicted to be highest directly off the coast of Tampa, Florida (Fig. 6G). This was caused by a likely bloom event of the mixotrophic dinoflagellate, *Karenia brevis*, captured in our DNA samples (fig. S11) and confirmed to be highly abundant through manual counts (10^5^–10^6^ cells l^−1^) estimated around the same time and location via the Florida Fish and Wildlife Conservation Commission (https://myfwc.com/research/redtide/monitoring). HABs formed by *K. brevis* are common in the GOM in the summer–fall, particularly along the West Florida Shelf, and can negatively impact marine ecosystems and local economies (*32*, *33*, *35*). There is evidence that warming may increase toxin production, growth rates, bloom frequency, and range expansion of some HAB species (*88*). Temperature had negative effects on Dinophyceae in our models (Fig. 5A–B), but responses were not explored to genus level. In culture, *K. brevis* has shown increased growth rates with increasing *p*CO_2_, though changes in toxin production were not recorded (*89*). It remains important to monitor HABs and their drivers (*90*), combining traditional monitoring and molecular methods to better predict pervasive blooms in the GOM and elsewhere.

### Indicator analysis reveals candidate microbial indicator taxa of OA

It is also important to determine specific microorganisms below the order to class level that may be indicative of different OA conditions in natural waters (*9*, *10*, *23*). To this end, we grouped samples in the photic zone (Cluster 1) based on TA:DIC ratios and examined microbial indicator taxa at the ASV level. The TA:DIC ratio is a well-used proxy for carbonate chemistry in the ocean, determining the buffering capacity against acidification (*79*, *91*). In general, lower TA:DIC ratios indicate poorly buffered waters, and so in our case, microbes that were more prevalent in lower TA:DIC samples may be candidate indicators of more acidic conditions in the GOM. TA:DIC ratios ranged from 1.1–1.2 in the photic zone, were not influenced by sampling transect, and were positively correlated with pH (Pearson *R* = 0.71; *P* < 0.01) in surface waters (Fig. 7A–B). TA:DIC ratios were manually grouped into low (< 1.16) vs. high (> 1.16) categories to explore microbial indicators (Fig. 7A–B).

**Figure 7:**
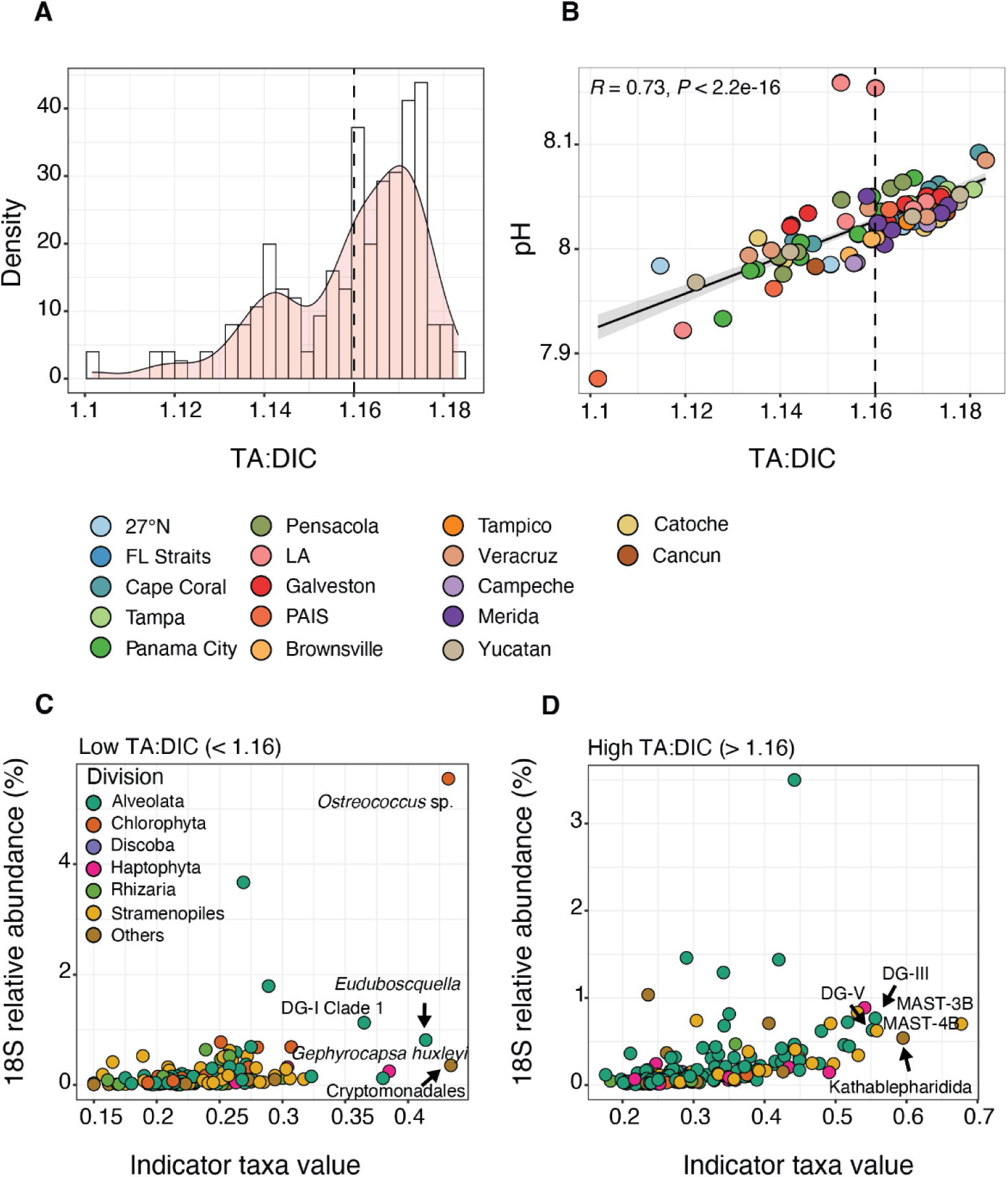
Protist indicator taxa based on TA:DIC ratios in the photic zone. **(A)** Histogram showing the density distribution of 18S samples in the photic zone (Cluster 1) based on TA:DIC ratios. **(B)** Values of in situ pH vs. TA:DIC in the photic zone, with samples colored by transect. Pearson correlation between variables is shown, with 95% confidence interval. The dotted line in panels **A**–**B** indicate the manual cutoff used for indicator analysis: low TA:DIC < 1.16 vs. high TA:DIC > 1.16. **(C**–**D)** Indicator values vs. average relative abundance (%) for protist ASVs in the photic zone that were significant to the analysis (*P* < 0.001) in samples with either low TA:DIC **(C)** or high TA:DIC **(D)**. Protist ASVs are colored by division and the top five ASVs with the highest indicator values are labeled in each panel, identified to their lowest possible taxonomic assignment (via the PR2 database). DG = Dino-Group. See table S2 for a full list of 18S (and 16S) indicator ASVs. Similar plots for 16S ASVs are shown in fig. S12.

Overall, we found that 146 and 117 protist ASVs were significant indicators (*P* < 0.001) of low or high TA:DIC ratios, respectively (table S2). Protist indicators spanned a range of taxonomic groups, though several ASVs stood out (Fig. 7C). For instance, protists with the highest indicator values (> 0.35) in samples with low TA:DIC ratios included *Ostreococcus* sp., which was the most relatively abundant indicator ASV on average in the photic zone (∼6%), as well as other ASVs assigned to *Emiliania huxleyi* (now *Gephyrocapsa huxleyi*), Cryptomonadales, *Euduboscquella* (Syndiniales), and Dino-Group I Clade 1 (Syndiniales). In comparison, ASVs with high indicator values (> 0.55) in samples with high TA:DIC ratios consisted of ASVs assigned to heterotrophic flagellates like MAST 3-B (and 4-B) and Kathablepharidida, as well as parasites in Dino-Group III and V (Fig. 7D). For 16S samples, a total of 228 and 136 ASVs were significant indicators (*P* < 0.001) of low vs. high TA:DIC ratios (table S2), dominated by Proteobacteria (fig. S12). 16S ASVs with the highest indicator values (> 0.45) in low TA:DIC samples included an ASV assigned to SAR11 clade Ia, which accounted for 10% of reads on average in Cluster 1, as well as other ASVs assigned to SAR11 (clade I and Ia), SAR406, and AEGEAN-169 (fig. S12). The 16S ASVs that were most indicative of high TA:DIC were assigned to SAR116, SAR86, AEGEAN-169, and Rickettsiales (family S25-593; fig. S12).

Pico- and nanoeukaryotes dominate warm and oligotrophic regions like the open GOM (*27*, *53*, *64*) and are sensitive to changing ocean conditions (*15*, *23*). Two of arguably the most well studied taxa in the field of phytoplankton OA research, *Ostreococcus* sp. and *Emiliania huxleyi*, were associated with less buffered (and more acidic) waters in the GOM (Fig. 7C). Both species are widespread and impact global biogeochemical cycles (*92*), with *E. huxleyi* being a major calcifier and contributor to CaCO_3_ flux (*93*). In a prior 18S rRNA metabarcoding survey in the southern GOM, *Ostreococcus* was the only genus with significantly different relative abundance between upwelling and downwelling conditions in the DCM and when comparing the DCM to mixed layer (*38*), which authors suggest may make this species an indicator of vertical nitrate flux. Our findings imply *Ostreococcus* may also be a candidate indicator of acidic conditions in GOM surface waters. Calcifying plankton, like *E. huxleyi*, are thought to be strongly impacted by OA, with increased *p*CO_2_ and/or lower pH having detrimental effects on growth and calcification rates (*15*, *16*). However, contrasting effects have been observed and may reflect considerable strain and ecotype variability (*18*, *23*). Indeed, several culture-based studies with *E. huxleyi* (and *Ostreococcu*s) have revealed adaptive mechanisms of cells to elevated *p*CO_2_ over hundreds of generations (*94*, *95*). Though *E. huxleyi* was not prevalent overall in our samples (fig. S8), this species has been measured in high concentrations (∼10^4^ cells l^−1^) in the southern GOM in spring (96). Together with model results at the class level (positive temperature effects on Prymnesiophyceae), our findings highlight the potential sensitivity of haptophytes to changing conditions in the GOM that should be further explored.

Indicator analysis also revealed SAR11, specifically ASVs assigned to clades 1 and 1a, as being possible indicators of less buffered waters in the GOM in summer–fall (fig. S12). SAR11 is the most abundant bacterial group in the oceans, playing an important role in global carbon cycling Though diverse, members of the SAR11 clade Ia ecotype tend to be most prevalent in surface oceans (*52*), adapted to nutrient-poor conditions via small cell sizes and streamlined genomes (*98*). Though direct effects of OA on SAR11 remain unclear and are likely to be less important compared to shifts in DOM (*21*), SAR11 exhibits known seasonality in the surface oceans and is sensitive to temperature (*48*, *65*). Such temperature sensitivity was supported in our model analysis (Fig. 4A–B). Given the ubiquity of SAR11 and its role in global carbon cycles, it remains critical to confirm and further investigate the potential of this group as an indicator of ocean change.

### Future sampling to characterize microbes in changing oceans

Efforts to characterize microbial communities over natural physicochemical gradients are essential to inform how these communities may shift in the face of changing ocean conditions (*10*). In the GOM, there is evidence of increased *p*CO_2_ in many parts of the open ocean that are on par with rates of change in other oligotrophic regions (*99*), like those measured in the Pacific Ocean via the Hawaiian Ocean Time-series (1.72 µatm yr^−1^) and in the Atlantic Ocean via the Bermuda Atlantic Time-series Study (1.69 µatm yr^−1^). Yet, knowledge on the effects of OA and warming on biological organisms is limited in the GOM, particularly for microbes. Here, we performed the first basin-scale DNA metabarcoding survey in the GOM and paired this with extensive hydrographic, nutrient, and carbonate chemistry measurements to investigate diverse prokaryotes and protists and their specific environmental drivers (Fig. 1). In line with prior physiological and modeling-based observations, our GOM model analyses suggest that more acidic and warmer conditions in the GOM may favor heterotrophic bacteria (SAR11 and SAR86) and smaller phytoplankton (e.g., Prymnesiophyceae), with groups like Dinophyceae potentially being less favored in future conditions (Fig. 4; Fig. 5). Warming and OA in the GOM may have contrasting effects on major plankton parasites (Syndiniales) and grazers (Sagenista) that are seldom considered with respect to climate change and underscores the importance to measure multiple stressors simultaneously (Fig. 5). We also defined microbial indicator taxa at the ASV level (Fig. 7), which resulted in several ubiquitous (and environmentally sensitive) microbes, like *Ostreococcus* sp., *Emiliania huxleyi*, and SAR11 clade Ia, being associated with more acidic waters in the GOM. Model inference and the utility of identified ASVs to act as indicator species of OA will need to be further tested, including at different times of the year to reflect seasonal turnover of the microbial community.

Though still unclear, empirical and predictive work suggests that changes in our ocean systems will likely have profound impacts on microbial composition, biogeography, and physiology (*8*, *100*), with consequences for trophic transfer, nutrient cycling, and carbon export. Global models and experimental evidence predict increased stratification with warming, shifting communities to smaller picophytoplankton that can better exploit nutrients and other resources (*12*, *14*). Warming-induced stratification may also result in an overall net reduction in carbon export that may threaten to decrease the amount of organic carbon that reaches the seafloor (*101*). Yet, predicted shifts in carbon export in global ecosystem models remains uncertain, ranging from a 41% decrease to 8% increase in carbon export flux in future oceans (*102*). Strong selection imposed by climate change may also drive rapid adaptation, competition, or the emergence of new species (e.g., with higher thermal tolerance), all restructuring microbial communities (*8*, *13*, *85*). In culture, some microbes demonstrate the ability to adapt to warmer or more acidic conditions (*94*, *95*), though this does not necessarily mean they will remain competitive and it remains an open question on how this will apply to natural systems with mixed assemblages (*19*, *85*). Further, current models do not fully account for trophic interactions, like grazing or parasitism, the rates of which will likely vary in future oceans and offset direct physiological effects of OA or warming on certain microbes. It remains important to measure microbial interactions, plankton mortality rates, and carbon export rates over time and in space (*10*, *102*), which will support a more mechanistic approach to model predictions.

Our findings provide an important baseline for microbial OA research in the GOM; however, sampling on GOMECC-4 only reflected a single time of the year and did not consider known seasonal variability in carbonate chemistry parameters or hydrography (*79*, *80*), which are likely to influence microbes and their drivers (*36*). In response, there is a need for sustained biological measurements in the GOM, either by establishing long-term monitoring programs or continuing to leverage existing oceanographic surveys, like GOMECC. Long term microbial sampling in the GOM will be essential to accurately predict future changes in microbial groups that may be expected with continued OA or warming. For example, increased DNA collection would support ecosystem modeling of microbes in the GOM, integrating climate model scenarios (e.g., via the Coupled Model Intercomparison Project) to predict shifts in microbial abundance by the end of the century. Ultimately, our ability to predict the response of marine microbes to climate change will depend on sustained and coordinated sampling efforts across a range of dynamic marine ecosystems.

## Materials and Methods

### Seawater collection, DNA filtration, and environmental metadata

Seawater was collected on board the NOAA Ship *Ronald H. Brown* as part of GOMECC-4 from September 13 to October 21, 2021. Sampling for GOMECC-4 occurred along 16 inshore– offshore transects across the entire GOM and an additional line at 27°N latitude in the Atlantic Ocean (Fig. 1A). Sampling started at the 27°N line and continued counterclockwise across the GOM, ending at Florida Straits and Cape Coral. We also collected DNA samples near Padre Island National Seashore (U.S. National Parks Service), a barrier island located off the coast of south Texas (Fig. 1A). Vertical CTD sampling was employed at each site to measure discrete physical, chemical, and biological properties. Water sampling for DNA filtration was conducted at 51 out of 141 total sites and three depths per site, representing the surface, deep chlorophyll maximum (DCM), and near bottom (fig. S1).

At each respective site and depth, seawater was collected from pre-designated Niskin bottles on a CTD rosette. To ensure adequate amounts of water were filtered for DNA analysis, samples for chemical parameters were taken at the same depths but with different discrete Niskin bottles. Following a CTD cast, which varied in duration from 30 min to 3 h depending on water depth, whole seawater was transferred from Niskin bottles to triplicate Whirl-Pak bags (3 depths x 3 replicates = 9 bags per site). Within an hour, whole seawater (∼2 L per replicate) was filtered through 0.22-µm Sterivex filters (Millipore; CAT# SVGP01050) via a peristaltic pump (100–150 rpm) and run dry. Filters were capped and outlets were sealed with parafilm. Filters were stored at –80 °C on the ship and kept at the same temperature in the laboratory for longer-term storage. Filter lines were sterilized with 2% bleach and rinsed with Milli-Q after every site. Milli-Q blanks were also filtered randomly throughout the duration of the cruise. Accounting for blanks and replication, a total of 481 Sterivex filters were collected on GOMECC-4.

Discrete samples for water column hydrography and chemistry were taken at each site and depth during GOMECC-4, including sites sampled for DNA. Temperature, salinity, pressure, and chlorophyll fluorescence were obtained from the CTD. Vertical CTD profiles on the downcast were used to estimate the position of the DCM at each site. Blanks and quality control samples were considered for each discrete chemical parameter. Dissolved oxygen concentration was estimated from water samples (125 ml) using an automated oxygen titrator with amperometric end-point detection (*103*). Nutrient samples were collected from Niskin bottles into 50-ml acid washed bottles. Dissolved nutrients (NO_3_, NO_2_, NH_4_, PO_4_, and SiO_4_) were measured on board using an automated continuous flow analytical system with colorimetric detection ((*104*); SEAL Analytical). Samples for DIC were collected from Niskin bottles into 294-ml borosilicate glass bottles, sealed with glass stoppers, and stored for 12 h at room temperature. DIC samples were analyzed on the ship using two analytical systems, each consisting of a coulometer (CM5017, UIC Inc.) coupled with a Dissolved Inorganic Carbon Extractor (*105*).

Samples for total alkalinity (TA) were collected from Niskin bottles into 500-ml collection bottles, preserved with a mercuric chloride solution, and kept in a water bath at 22 °C for 1 h prior to analysis. TA measurements were made using a two-titration system, consisting of a Metrohm 765 or 665 Dosimat Titrator and Orion 720A or 2-Star pH meter (*106*). Samples for *p*CO_2_ were drawn from Niskin bottles into 500-ml glass bottles, preserved with mercuric chloride, and stored at room temperature for 8 h before analysis. Details on the system used to measure *p*CO_2_ are described in (*107*) and include equilibrating each sample with a constantly circulating gas phase. Lastly, for pH analysis, samples were collected from Niskin bottles into 10-cm (∼30 ml) glass cylindrical optical cells and analyzed on an Agilent 8453 spectrophotometer with a custom-made temperature-controlled cell holder (*108*). Aragonite saturation state was calculated at each site and depth based on temperature, salinity, pressure, DIC, and TA using the CO2SYS program for CO_2_ System Calculations (*109*). Measurements of *p*CO_2_ (20 °C) and pH (25 °C) were re-calculated to in situ conditions using pressure, temperature, salinity, DIC, and TA in CO2SYS (*109*). Environmental metadata associated with DNA samples are provided in table S3.

### DNA extractions, PCRs, and library preparations

Sterivex filters were extracted in-house at NOAA’s Atlantic Oceanographic and Meteorological Laboratory (AOML) using the ZymoBIOMICS 96 DNA/RNA MagBead kit (Zymo; CAT# D4308), with modifications for in-cartridge bead beating as described in (*110*). Filters were thawed, the inlet caps were removed, and excess water was dried from the inlet using kimwipes to allow for dispensing of beads into the cartridge. Premade mixtures of 0.1 mm and 0.5 mm beads were directly added into the filters to ensure adequate lysis and recovery of hard-to-lyse phytoplankton groups (*110*). This was followed by the addition of a lysis buffer (1 ml). Sterivex filters were vortexed for 40 min on a Vortex-Genie at maximum speed (∼3200 rpm). DNA lysates were transferred to 2-ml LoBind tubes (Eppendorf) via syringe and centrifuged for 1 min at 10,000 g. Supernatant (750 µl per sample) was split across three KingFisher 96-well plates (250 µl per plate). Zymo MagBinding buffer (600 µl) and magnetic beads (25 µl) were added to each well in each of the three plates. With this setup, 96 samples were extracted at the same time on the automated KingFisher Flex (Thermo Fisher). Each run included three wash plates with 500–900 µl per well of MagWash and an elution plate with 150 µl per well of molecular-grade water. DNA was eluted into a single well from the same discrete sample across replicate plates. Concentrations of eluted DNA were measured using a Varioskan LUX plate reader and the Quant-IT dsDNA Assay (Thermo Fisher) and corrected per replicate sample based on volume of seawater filtered (ng l^−1^; fig. S13). Filters were processed randomly. Extraction blanks (clean Sterivex filters) were also included and processed similarly. A bacterial mock community (Zymo) was included as a positive control.

Metabarcoding libraries were initially prepared at AOML, amplifying DNA of target organisms with universal primers, including 16S (Bacteria and Archaea) and 18S rRNA (protists). Primers from (*111*) were used to target the 16S V4–V5 region: forward (515f; 5’-GTGYCAGCMGCCGCGGTAA-3’) and reverse (926r; 5’-CCGYCAATTYMTTTRAGTTT-3’). Primers from (*112*) and the Earth Microbiome Project (http://www.earthmicrobiome.org/emp-standard-protocols/18s/) targeted the 18S V9 region: forward (1391f; 5’-GTACACACCGCCCGTC-3’) and reverse (EukBr; 5’-TGATCCTTCTGCAGGTTCACCTAC-3’). Primers were constructed with Fluidigm common oligos CS1 forward (CS1-TS-F: 5’-ACACTGACGACATGGTTCTACA-3’) and CS2 reverse (CS2-TS-R: 5’-TACGGTAGCAGAGACTTGGTCT-3’) fused to their 5’ ends, to enable two-step library preparation at the Michigan State University Research Technology Support Facility (RTSF).

PCR reactions were run in triplicate (12.5 µl per sample), with 1 µl of DNA per sample. 16S PCR reactions consisted of 5 µl of AmpliTaq Gold, 6.25 µl of water, and 0.375 µl of each primer (10 µM); PCR conditions included denaturation at 95 °C for 2 min, 25 cycles of 95 °C for 45 s, 50 °C for 45 s, and 68 °C for 90 s, followed by a final elongation step of 68 °C for 5 min (*111*). 18S PCR reactions consisted of 5 µl of AmpliTaq Gold, 6.5 µl of water, and 0.25 µl of each primer (10 µM); PCR reactions involved denaturation at 94 °C for 3 min, 35 cycles of 94 °C for 45 s, 65 °C for 15 s, 57 °C for 30 s, and 72 °C for 90 s, followed by a final elongation step of 72 °C for 10 min (*112*). PCR products were pooled and run on a 2% agarose gel to confirm amplification of target bands. Sample plates were submitted to the Michigan State University RTSF Genomics Core for secondary PCR and sequencing.

Secondary PCR used dual-indexed, Illumina-compatible primers, targeting the Fluidigm CS1/CS2 oligomers at the ends of the PCR products. PCR conditions for the secondary run included an initial denaturation step at 95 °C for 3 min, 11 cycles of 95 °C for 15 s, 60 °C for 30 s, and 72 °C for 60 s, followed by elongation at 72 °C for 3 min. Amplicons were batch normalized using Invitrogen SequalPrep DNA Normalization plates and the recovered product was pooled. The pool was QC’d and quantified using a combination of Qubit dsDNA HS, Agilent 4200 TapeStation HS DNA1000, and Invitrogen Collibri Library Quantification qPCR assays. The RTSF Core included a sequencing blank for each sample plate. Separate sequencing runs were performed using an Illumina MiSeq (2 × 250 bp) for 18S and 16S samples. Custom sequencing and index primers complementary to the Fluidigm CS1 and CS2 oligomers were added to appropriate wells of the reagent cartridge. Base calling was done by Illumina Real Time Analysis (RTA) v1.18.54 and output of RTA was demultiplexed and converted to FASTQ format with Illumina Bcl2fastq v2.20.0.

### Bioinformatics and functional assignments

Primers were removed from demultiplexed FASTQ sequences using Cutadapt (*113*). Trimmed reads were processed in Tourmaline, which implements QIIME 2 (and DADA2 plugins) in a Snakemake workflow (*114*). Paired-end DADA2 was used to infer 16S and 18S amplicon sequence variants or ASVs (*115*). Taxonomic assignments were also performed in Tourmaline using reference files from SILVA (Version 138.1; (*116*)) and the Protistan Ribosomal Reference or PR2 (Version 5.0.1; (*117*)) databases for 16S and 18S ASVs, respectively. In both cases, taxonomy was assigned using a Naïve Bayes classifier trained to the respective databases and trimmed to the primer regions (*118*). Output files (taxonomy, count, and metadata) were imported separately into R (Version 4.3.1) using qiime2R (Version 0.99.6; https://github.com/jbisanz/qiime2R) and merged with phyloseq (Version 1.44.0; (*119*)). Several groups were removed from the 18S dataset: Metazoa, Streptophyta, Rhodophyta, and unassigned reads at the Subdivision level. 18S reads assigned to non-marine taxa, e.g., Insecta, Archosauria, and Ascomycota were also filtered out. For 16S, reads assigned to Chloroplast, Mitochondria, and Eukaryota were removed. Samples with less than 3,000 reads counts were filtered out for 18S (5,000 reads for 16S), along with ASVs only observed once in each respective dataset. Species accumulation curves were generated for 18S and 16S samples using the R package ranacapa (Version 0.1.0; (*120*)). The number of reads vs. ASVs was saturated with respect to categorial depth and position of samples on the shelf vs. open GOM, indicating that an appropriate sequencing depth was reached (fig. S14). Samples were rarefied to the minimum read count to normalize for differences in library size.

Protist ASVs were manually assigned to functional groups based on 18S V9 functional annotations (https://doi.org/10.5281/zenodo.3768950) that were previously applied to Tara Ocean communities (*57*). Additional databases (e.g., World Register of Marine Species) and literature searches were also used. The following functional groups were included for 18S protists: autotrophic protists, heterotrophic protists, mixotrophic protists, parasites, photosymbionts, and other protists. Mixotrophic protists were further categorized as being constitutive mixotrophs (CM) that inherently have chloroplasts and endosymbiotic specialist non-constitutive mixotrophs (eSNCM) that harbor endosymbionts to support growth (*121*). We recognize that many protists likely exhibit mixotrophy in some capacity, and so, our functional annotation of this group may be underrepresented. Other protists represented higher level taxonomic groups (domain or supergroup) that were unassigned at lower levels. Bacteria and Archaea were categorized functionally as being heterotrophic or autotrophic.

### Statistical analyses

Prior to ordination, ASV count tables were transformed to Aitchison distances, which is estimated by transforming read counts via centered log-ratio normalization and computing Euclidean distances. The resulting Aitchison distance matrices were used to observe microbial composition and aimed to minimize compositional bias inherent with amplicon data (*83*). Principal coordinate analysis (PCoA) of Aitchison distances was used to visualize 16S and 18S community composition. Permutational multivariate analysis of variance (PERMANOVA) tests were employed with the adonis2 function in vegan (9999 permutations) to estimate the impact of spatial factors on community composition. This included categorical depth (surface, DCM, and near bottom), sampling transect, and location of samples on the continental shelf vs. in open ocean regions of the GOM designated by the 200 m isobath (Fig. 1A).

Samples were also grouped into clusters via hierarchical clustering (Ward’s method) based on Aitchison distances using the hclust function in vegan (Version 2.6-6.1; (*122*)). The optimal number of clusters was determined based on average silhouette widths using the factoextra package (Version 1.0.7; (*123*)). Silhouette widths offer an estimate on the quality of sample clustering, with higher width coefficients indicating optimal clustering (*124*). Three clusters were found to be optimal for both 16S and 18S (fig. S15), which largely reflected depth in the water column (Fig. 1B; fig. S1). Cluster 1 consisted of samples collected at all depths on the shelf and offshore in the surface layer, all confined to the photic zone (2–99 m). Cluster 2 consisted of samples mainly from offshore and more stratified waters in the DCM (2–124 m), while Cluster 3 represented samples collected offshore in meso-to bathypelagic waters (135–3,326 m). The photic zone extends to 200 m in many deeper regions of the GOM, and so, samples in Cluster 2 (and a handful in Cluster 3) were also technically collected within the photic zone. However, we distinguish communities in Clusters 2–3 from Cluster 1 based on the large proportion of samples confined to the open ocean DCM (Cluster 2; 80%) and mesopelagic (Cluster 3; 98%) that reflect disparate habitats in the GOM.

Mean Shannon diversity index and richness (# of ASVs) were determined for each cluster using the estimate_richness function in phyloseq (*119*) and compared against other clusters with Wilcoxon tests (*P* < 0.05). Mean diversity and richness were also estimated along transects, applying local regression (loess) curves to visualize trends using the geom_smooth function in ggplot2 (Version 3.5.1; (*125*)). Stacked bar plots displaying mean relative abundance were observed at the class level for 18S and order level for 16S for each sampling transect and cluster using the microeco package in R (Version 1.7.1; (*126*)). Taxonomic profiles were also observed using the treemap package in R (Version 2.4-4; (*127*)), a tiered approach to visualize relative abundance across multiple taxonomic levels.

Indicator taxa that were more abundant and representative of high (or low) TA:DIC ratios were statistically inferred using the indicspecies package in R (Version 1.7.14; (*128*)). The TA:DIC ratio was chosen because it is a good proxy to determine the ocean’s capacity to absorb anthropogenic CO_2_ by influencing its buffering capacity (*91*). Higher ratios indicate strong buffering capacity (i.e., the capacity of seawater to buffer against acidification). Based on histograms of TA:DIC, samples were grouped a priori into high (> 1.16) or low categories (< 1.16) that reflect different OA conditions (Fig. 7A–B). We focused on DNA samples collected from the photic zone (Cluster 1) to mitigate natural depth effects and to provide additional context to models (see next section). Indicator analysis was run separately on rarefied 16S or 18S samples that were agglomerated to the species level using the function multipatt with 999 permutations (*128*). Significant ASVs (*P* < 0.001) were retained and summarized for high (or low) TA:DIC and plotted against their mean relative abundance in the photic zone.

### Generalized linear models

Generalized linear models (GLMs) were used to examine relationships between environmental factors (predictor variables) and the relative abundance of major microbial groups (response variables). GLMs focused on DNA samples collected in the photic zone (Cluster 1), in large part to mitigate collinearity of factors that was prevalent in Clusters 2–3 (table S1). Separate GLMs were performed for the top four most relatively abundant order level 16S and class level 18S groups (Table 2). Separate models were constructed for *Synechococcus* and *Prochlorococcus* to resolve differences between major cyanobacteria genera. Only variables that met requirements of low collinearity (Spearman *r_s_* < 0.7 or > –0.7) and a variance inflation factor (VIF) < 10 were considered (*129*). Zurr et al. (2010) suggest using a more stringent VIF cutoff (< 3). However, we aimed to retain as many variables in the dataset as possible, which meant a few variables (e.g., DIC and salinity) approached VIF = 10. To select the best model for each 18S or 16S group, variables were further selected in a stepwise manner based on Akaike Information Criteria (AIC) values using the stepAIC function in the MASS package in R (Version 7.3-60; (*130*)). Only significant variables (*P* < 0.05) were used in the final model.

Final models were constructed with either Poisson or negative binomial error distributions. Initial model type was chosen by comparing standardized residuals and other model indices (e.g., AIC) with the compare_performance function in the performance package in R (Version 0.11.0; (*131*)). GLMs were implemented with the glm.nb function for negative binomial models in the MASS package or the glm function for Poisson models (family = Poisson) in the stats package in base R (Version 4.3.1). We observed overdispersion in relative abundance data for several Poisson models (Syndiniales, Dinophyceae, *Synechococcus*, and *Prochlorococcus*), in which case negative binomial models were applied (Table 2). Model quality and fit was estimated for each group using the check_model function in the performance package (*131*), which included plots of posterior predictive checks (model simulations), standardized residuals (Q–Q plots), homogeneity of variance, and collinearity of selected predictor variables. The goodness of fit was assessed with a pseudo *R*^2^ (Nagelkerke’s), though standardized residual checks of the final models were also carried out (*62*) to assess model fit and uniformity of the residuals (Kolmogorov–Smirnov, *P* > 0.05). As an additional validation, relative abundance data for each group was randomly split and trained with respective models using 80% of the data to predict a test set that was left out (20%). Pearson correlations were performed between model-trained and test data.

Model coefficients were scaled and visualized for each 18S and 16S group using the multiplot function in the coefplot package in R (Version 1.2.8; (*132*)). Individual factors were plotted against predicted relative abundance using the plot_model function in the sjPlot package in R (Version 2.8.16; (*133*)). We focused predictive plots on temperature and carbonate chemistry parameters (DIC and pH). Group-specific GLMs were used to predict relative abundance at all GOMECC-4 sites where surface layer (< 10 m) variables were collected (135 out of 141 sites). Six stations did not have representative CTD data available at the surface layer and were excluded. Predicted relative abundance for all surface GOMECC-4 sites were observed in Ocean Data View using Data-Interpolating Variational Analysis (DIVA) interpolation (*134*).

## Acknowledgements

We thank the captain and crew of the NOAA Ship *Ronald H. Brown* for logistical support for sample collection in the Gulf. We acknowledge the efforts of many scientists at NOAA and partnering institutions for collecting and processing oceanographic data on GOMECC-4. We thank Easton White and Elizabeth Harvey for their helpful conversations, and Sean McAllister for carefully reviewing the manuscript. We acknowledge the genomics services performed in the RTSF Genomics Core at Michigan State University.

## Author contributions

LRT, SRA, LB, and CRK conceived the study. SRA collected DNA samples. LB led collection of carbonate chemistry parameters. SRA and LRT processed DNA samples. SRA performed bioinformatics and data analysis. LRT, KS, and FAG contributed to improve data analysis. BAS and AS provided guidance on taxonomy and functional assignments. SRA and LRT led the writing of the manuscript. All authors contributed to revising the manuscript and approved the final version.

## Data and materials availability

Code and associated files needed to reproduce results and figures for this study are available on GitHub (https://github.com/aomlomics/gomecc) and have been archived on Zenodo (https://zenodo.org/records/13102580). All 18S and 16S sequence data generated from this study have been published at the National Center for Biotechnology Information (NCBI)’s Sequence Read Archive and BioSample database and are available with BioProject accession number PRJNA887898. Species count data generated from this study have been published on the Ocean Biodiversity Information System (OBIS) and the Global Biodiversity Information Facility (GBIF) at https://doi.org/10.15468/sm6fpz. Biological data has also been submitted to the National Centers for Environmental Information (NCEI) at https://www.ncei.noaa.gov/archive/accession/0250940/data/0-data/noaa-aoml-gomecc. Environmental measurements from the Niskin bottles and CTD profiles are also available at NCEI at https://doi.org/10.25921/4twf-pp50 and https://doi.org/10.25921/04h7-gv36, respectively. A cruise report detailing all the sampling and analyzing procedures during GOMECC-4 is available at https://doi.org/10.25923/rwx5-s713.

