## Supplementary Information for "Assessing the effects of warming and carbonate chemistry parameters on marine microbes in the Gulf of Mexico through basin-scale DNA metabarcoding"

Sean Anderson *et al.*

Corresponding authors: Sean Anderson and Luke Thompson


**This PDF file includes:**

Figs. S1 to S16
Legends for Tables S1 to S3

**Other Supplementary Materials for this manuscript include the following:**

Table S1
Table S2
Table S3

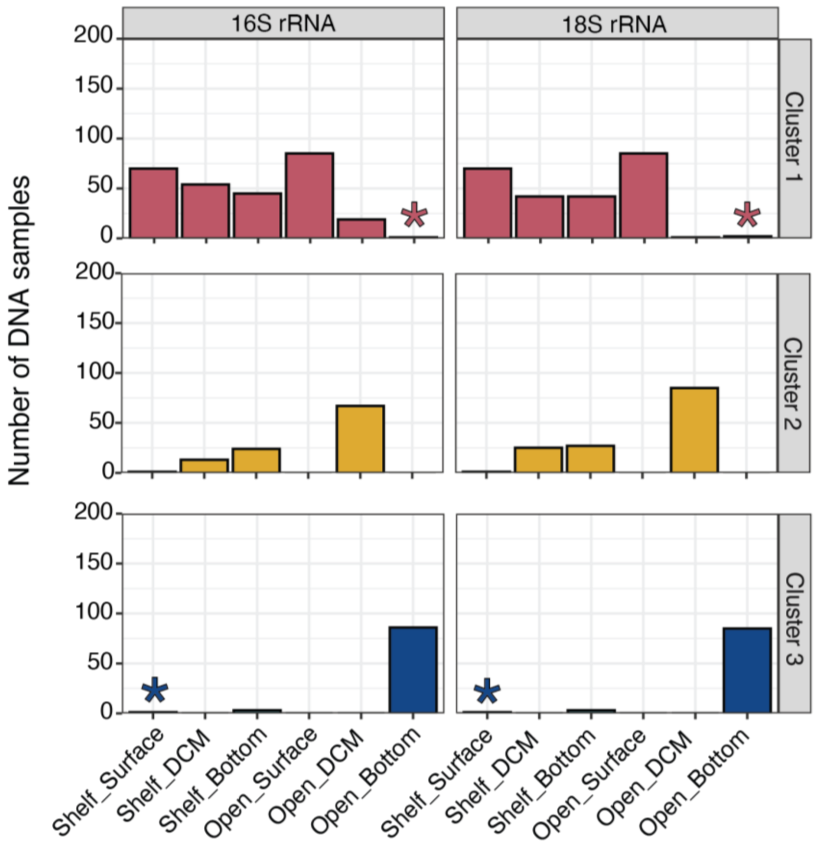


**Figure S1:** Hierarchical clustering was performed with amplicon data, which revealed three clusters (Clusters 1–3; top to bottom) for both 16S (left) and 18S (right) samples. Clusters were characterized by depth in the water column (surface, deep chlorophyll maximum or DCM, and near bottom) and position of samples on the continental shelf (< 200 m) vs. in open ocean regions of the GOM (> 200 m). Asterisks represent samples (three total; similar for 16S and 18S) that were removed from subsequent analysis due to their unexpected position in either Clusters 1 or 3.

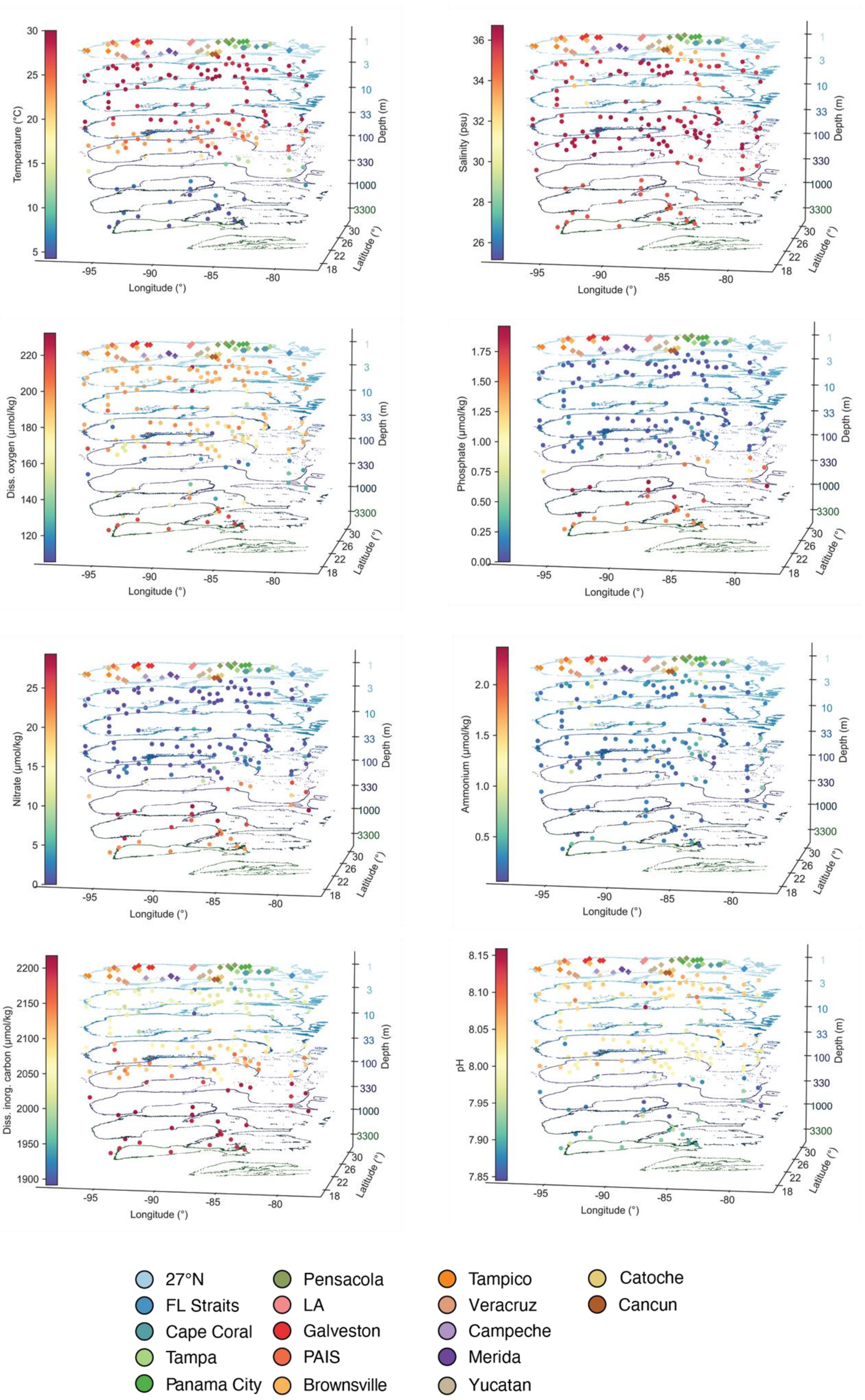

Temperature (°C)
Salinity (psu)
Diss. oxygen (µmol/kg)
Phosphate (µmol/kg)
Nitrate (µmol/kg)
Ammonium (µmol/kg)
Diss. inorg. carbon (µmol/kg)
pH
Depth (m)
1
3
10
33
100
330
1000
3300
Latitude (°)
30
26
22
18
Longitude (°)
-95
-90
-85
-80
27°N
FL Straits
Cape Coral
Tampa
Panama City
Pensacola
LA
Galveston
PAIS
Brownsville
Tampico
Veracruz
Campeche
Merida
Yucatan
Catoche
Cancun

**Figure S2:** Three-dimensional plots of depth-related (log scale) environmental factors that were considered for initial microbial GLMs (Table 1). Variables that were considered included temperature, salinity, oxygen ($O_2$), nitrate ($NO_3$), phosphate ($PO_4$), ammonium ($NH_4$), in situ (and recalculated) pH, and dissolved inorganic carbon (DIC). Stations are colored by transect at the surface. Spatial changes in values are shown for each parameter with a color gradient.

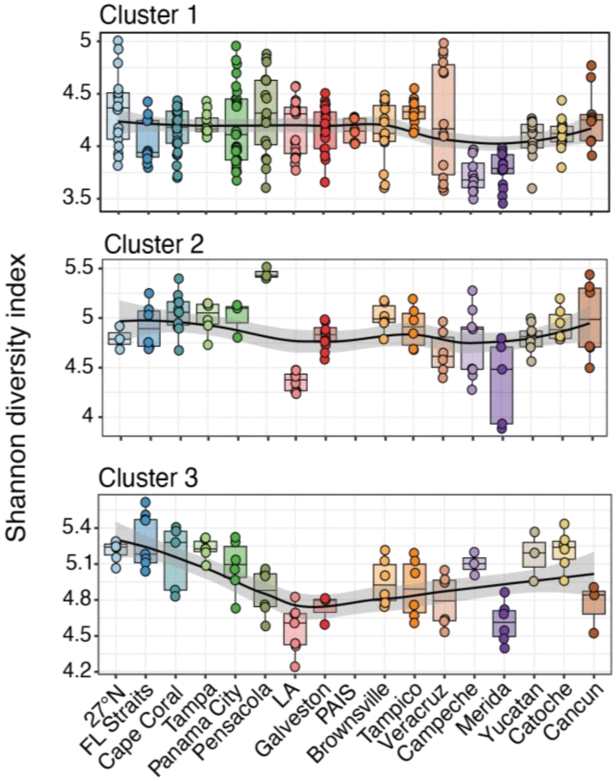

Cluster 1
Cluster 2
Cluster 3
Shannon diversity index
27°N
FL Straits
Cape Coral
Tampa
Panama City
Pensacola
LA
Galveston
PAIS
Brownsville
Tampico
Veracruz
Campeche
Merida
Yucatan
Catoche
Cancun


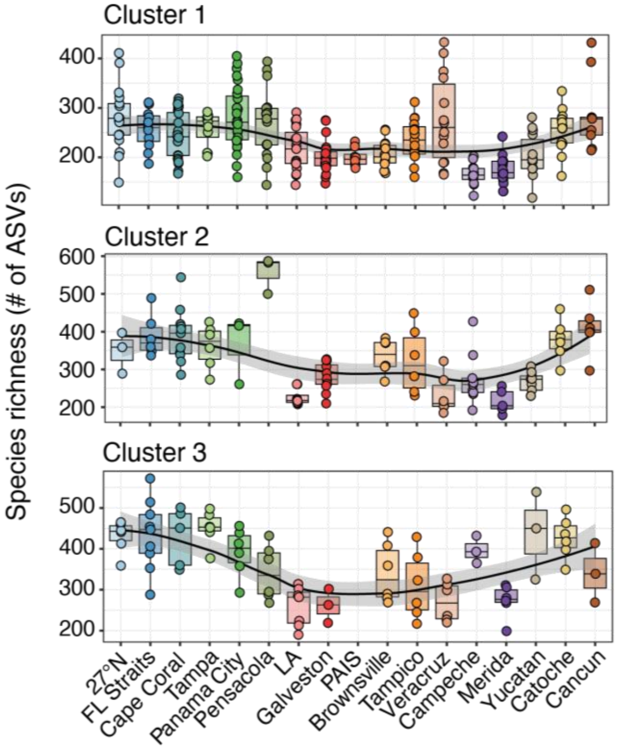

Cluster 1
Cluster 2
Cluster 3
Species richness (# of ASVs)
27°N
FL Straits
Cape Coral
Tampa
Panama City
Pensacola
LA
Galveston
PAIS
Brownsville
Tampico
Veracruz
Campeche
Merida
Yucatan
Catoche
Cancun

**Figure S3:** Mean Shannon diversity index (top) and species richness (# of ASVs; bottom) for 16S samples with respect to transect and faceted for each of the three clusters (Clusters 1–3). All data points are shown for each transect. Local regression (loess) curves were applied to the data and represent smoothed trends (black lines) with 95% confidence intervals. Sampling lines are ordered counterclockwise in the gulf according to the order of sampling; however, several lines (FL straits and Cape Coral) were sampled last but grouped here with other FL lines to more accurately visualize spatial patterns. LA = Louisiana and PAIS = Padre Island National Seashore.

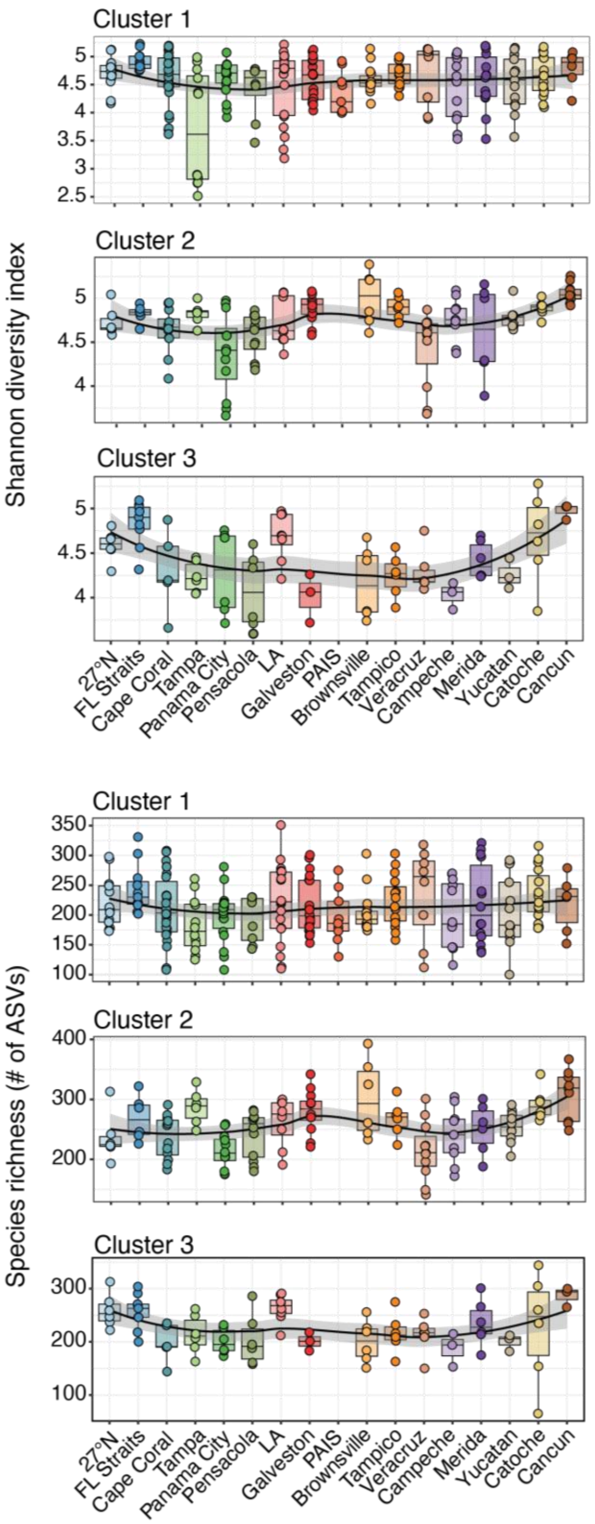


**Figure S4:** Same plot as in fig. S3 but for 18S samples, highlighting trends in Shannon diversity and species richness with respect to sampling transect and clusters.

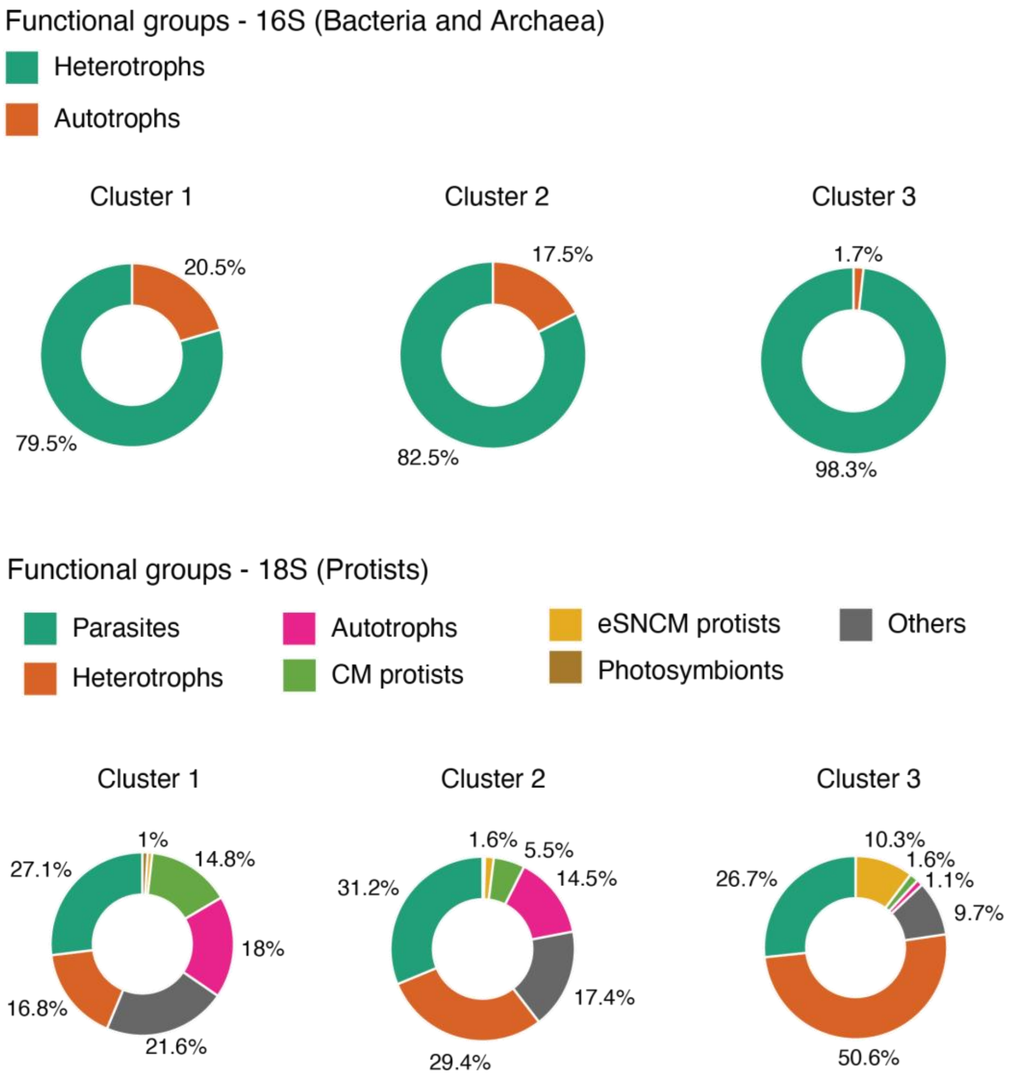


**Figure S5:** Percent of 16S (top) or 18S (bottom) ASVs (on average) that were manually assigned to different functional groups in Clusters 1–3. Groups were assigned manually to ASV tables based on prior functional databases and literature searches (see Materials and Methods). For 16S, groups included heterotrophs and autotrophs (cyanobacteria). Protists were categorized as being parasites, heterotrophs, autotrophs, constitutive mixotrophs (CM), endosymbiotic specialist non-constitutive mixotrophs (eSNCM), and photosymbionts. An "other protists" category was included and reflected taxa that were unassigned at the domain or supergroup according to the PR2 database.

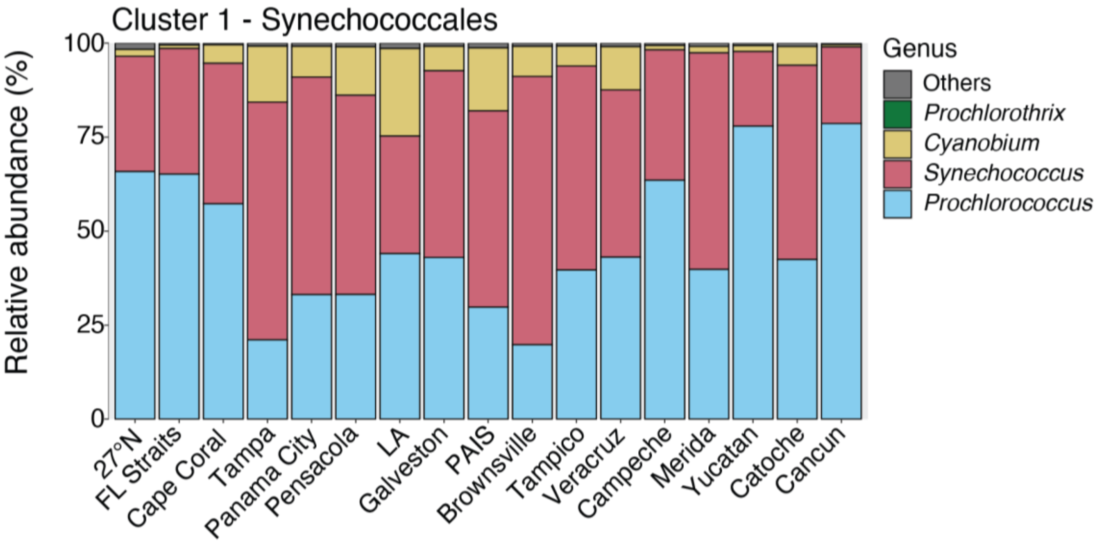


**Figure S6:** Stacked bar plots of mean relative abundance (%) at the genus level within the order Synechococcales in each transect in the photic zone (Cluster 1). Four genera dominated relative abundance (others in gray), including *Prochlorococcus*, *Synechococcus*, *Cyanobium*, and *Prochlorothrix*.

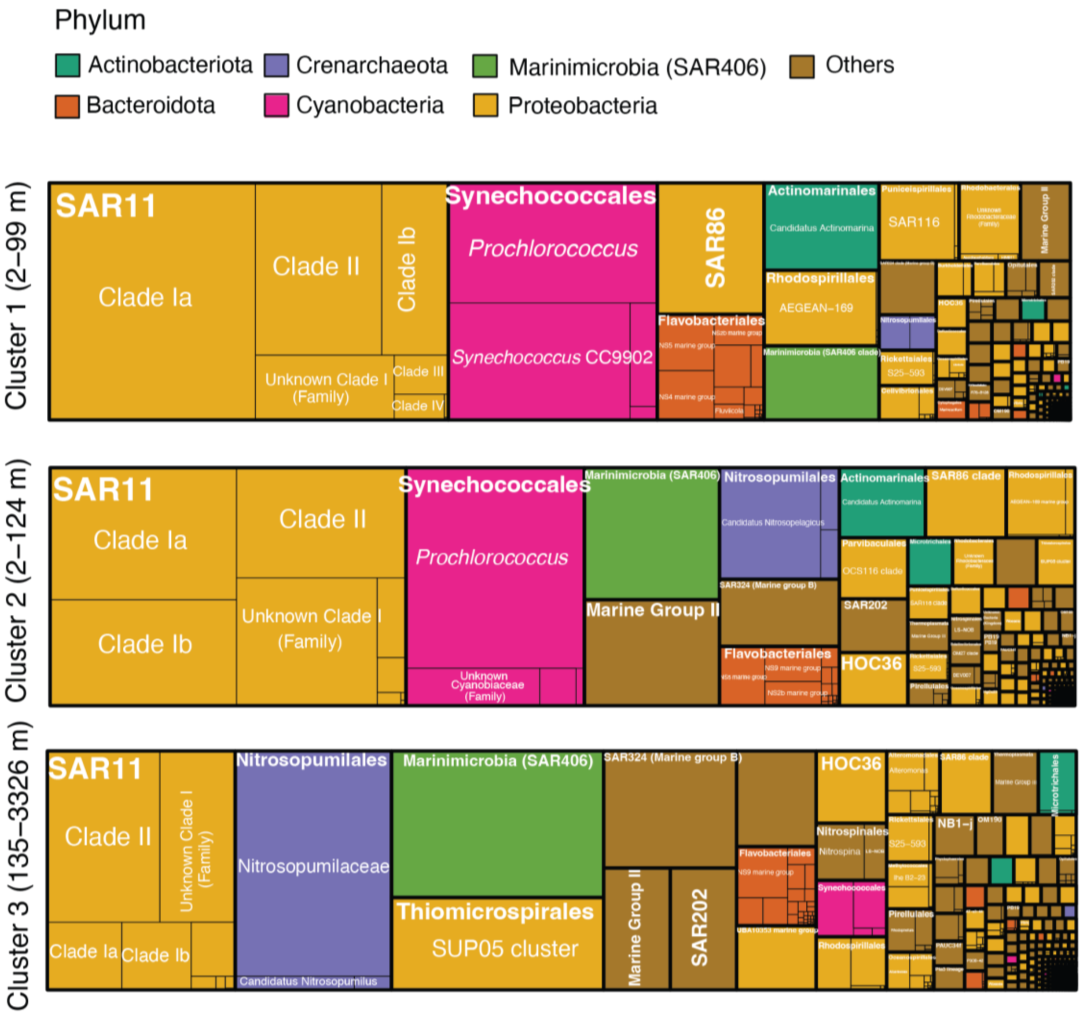


**Figure S7:** Taxonomy tree maps of 16S samples based on mean relative abundance (%) across all samples in Clusters 1–3. Taxonomic assignments were made via the SILVA database at the order level (larger boxes) and genus level within each order (smaller boxes). Boxes are colored by phylum and focused on the top six groups, with less abundant organisms at the phylum level grouped into an "others" category (in brown). Clusters largely reflected depth in the water column on the shelf vs. open ocean: Cluster 1 (photic zone; 2–99 m), Cluster 2 (DCM; 2–124 m), and Cluster 3 (aphotic zone; 135–3,326 m).

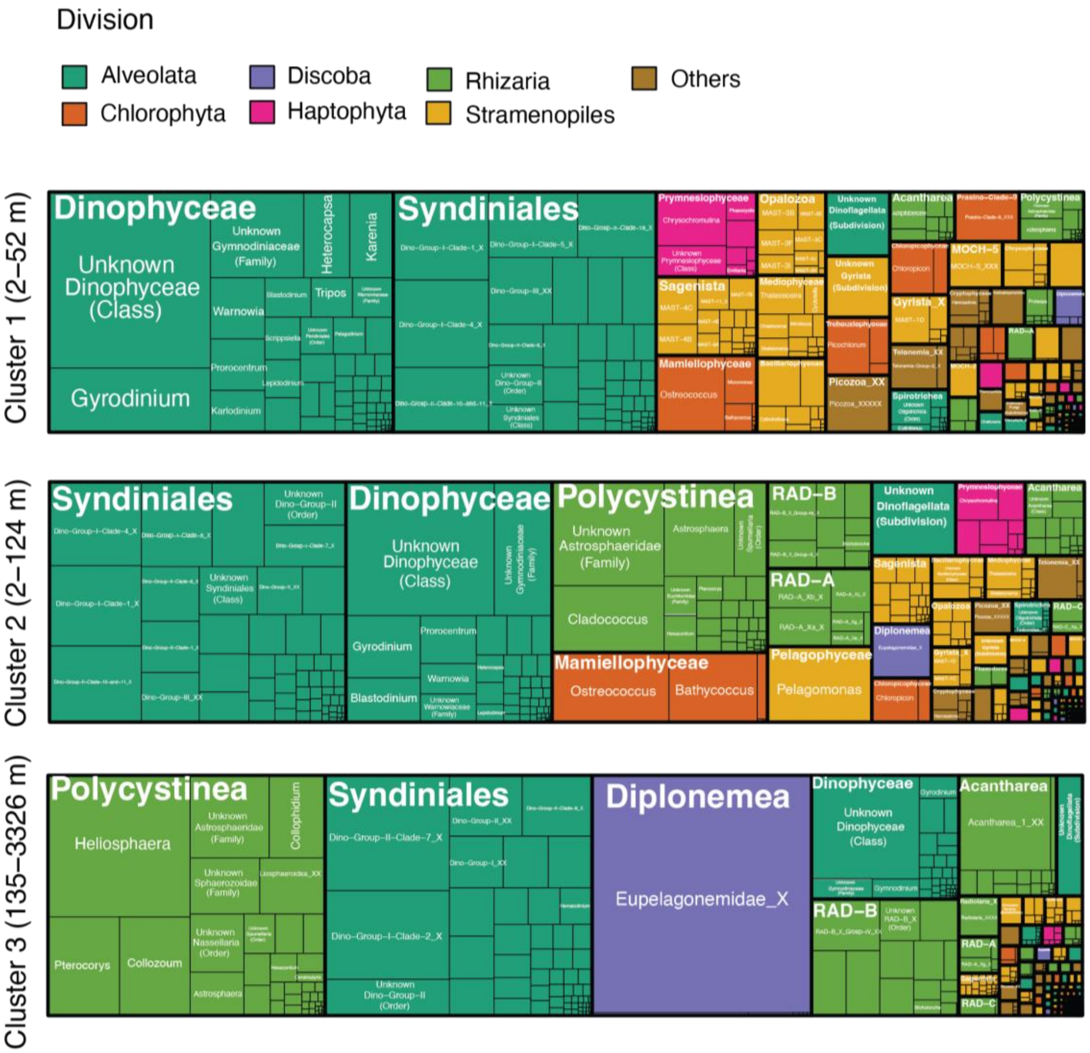


**Figure S8:** Taxonomy tree maps of 18S samples based on mean relative abundance (%) across all samples in Clusters 1–3. Taxonomic assignments were made via the PR2 database at the class level (larger boxes) and genus level within each class (smaller boxes). Boxes are colored by division and focused on the top six groups, with less abundant organisms at the division level grouped into an "others" category (in brown). Clusters were similar to those observed among 16S samples in fig. S7.

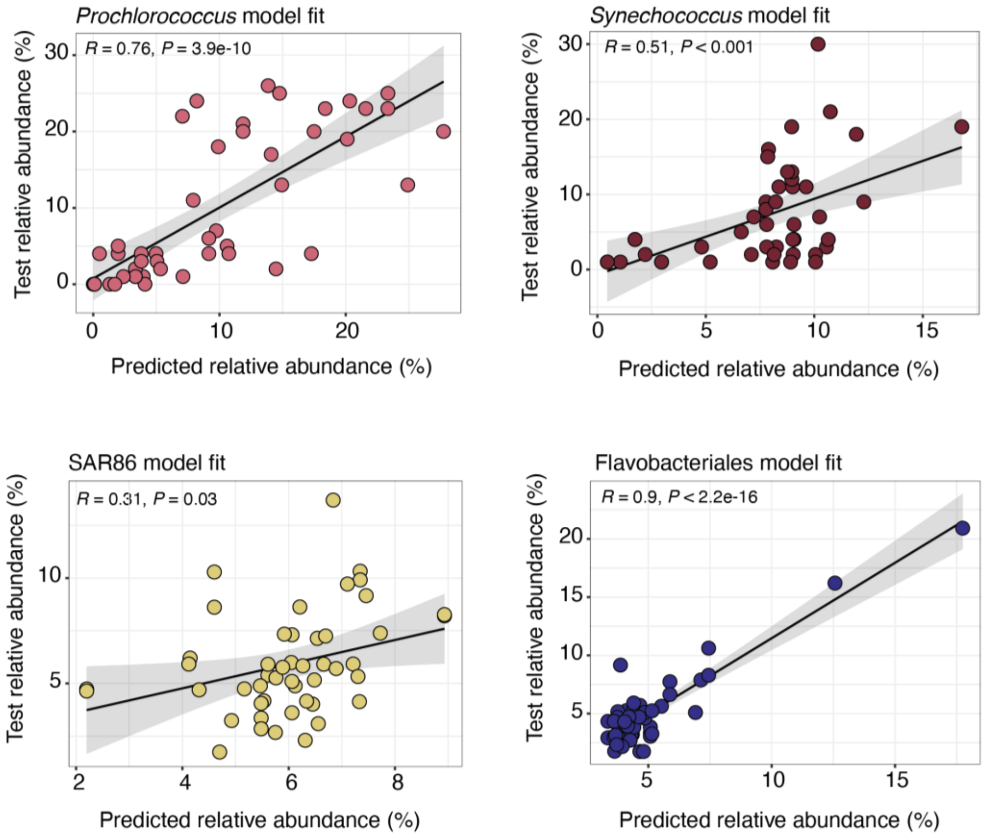


**Figure S9:** Testing the model fit of final GLMs for major 16S groups in the photic zone, which included *Prochlorococcus*, *Synechococcus*, SAR86, and Flavobacteriales. Model fit for SAR11 is shown in Fig. 4D. For each group, relative abundance data was split randomly (80:20) and GLMs were trained with 80% of the data, inclusive of triplicates (219 samples). Predicted relative abundance (%) derived from the trained set was correlated to test values that were held out (20%; 48 samples) via Pearson correlations. P-values are shown for each group and regression lines (with 95% confidence intervals) are also displayed.

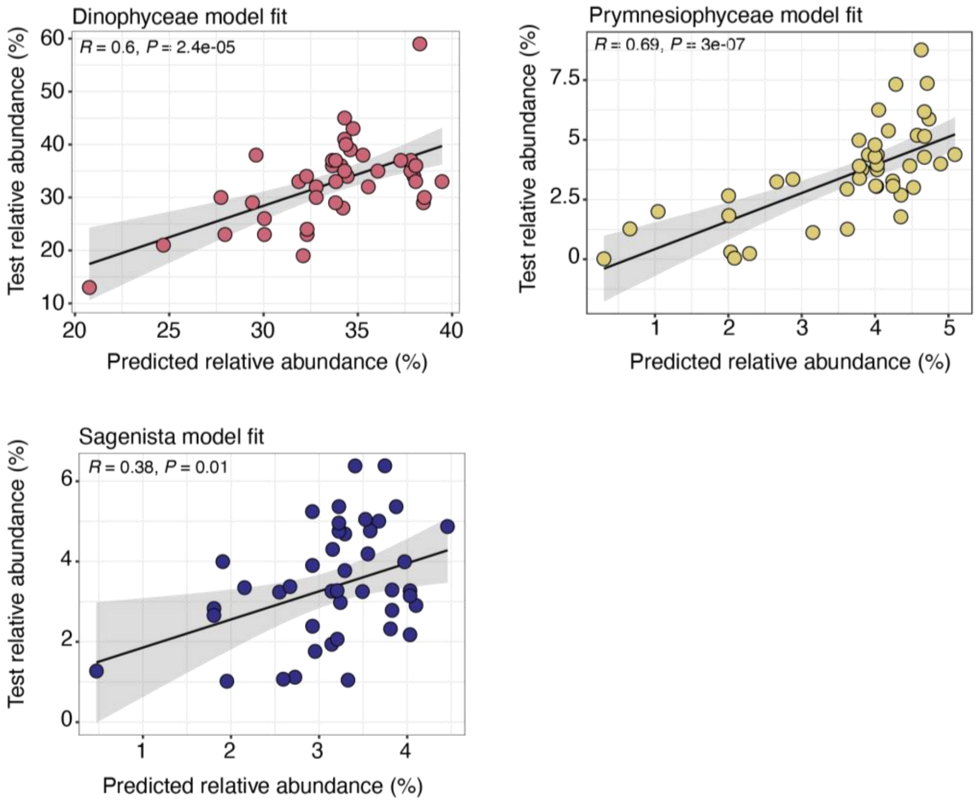


**Figure S10:** Testing model fit for major 18S groups in the photic zone, including Dinophyceae, Prymnesiophyceae, and Sagenista. Model fit for Syndiniales is shown in Fig. 5D. Other details are the same as in fig. S9 for 16S groups.

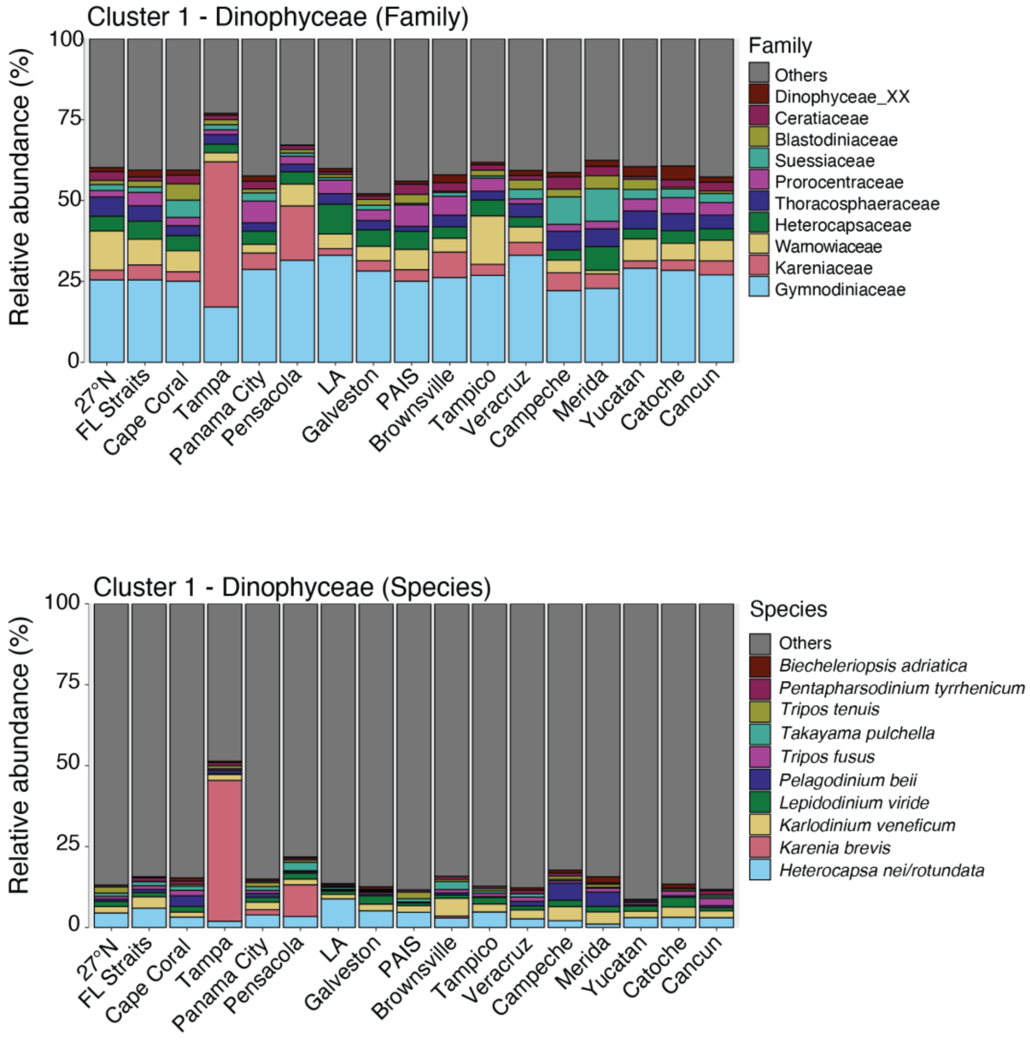


**Figure S11:** Stacked bar plots of mean relative abundance (%) for the top ten family (top panel) and species level (bottom panel) groups within Dinophyceae over sampling transects in the photic zone (Cluster 1). Less abundant taxa are marked as "others" (in gray). Taxonomic assignments were based on the PR2 database. Plots emphasize a likely harmful bloom of *Karenia brevis* (Kareniaceae) along the Tampa and Pensacola lines that corresponded to high concentrations (~$10^5$–$10^6$ cells $l^{-1}$) of *K. brevis* measured at the same time and location via the Florida Fish and Wildlife Conservation Commission.

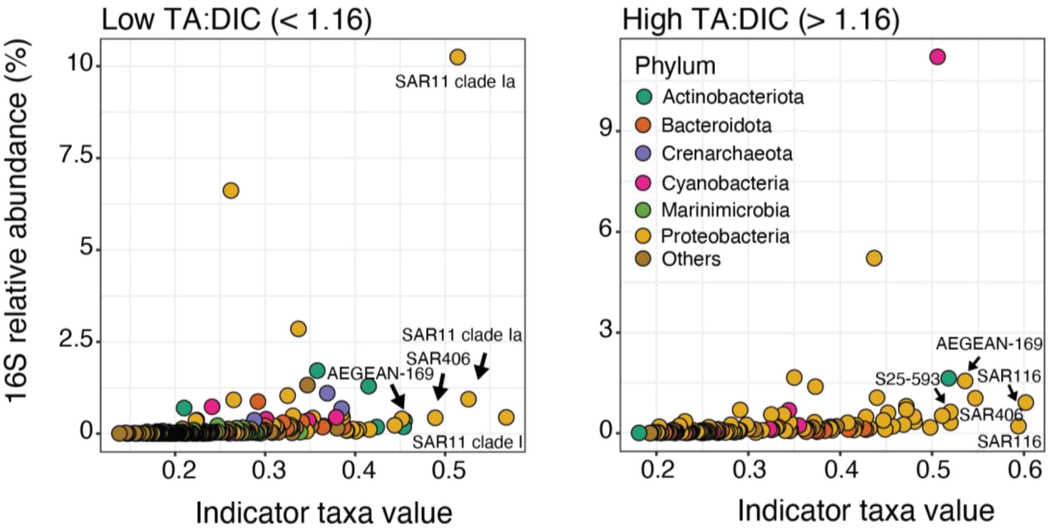


**Figure S12:** Microbial indicator values vs. mean relative abundance (%) of 16S ASVs that were significant indicators ($P < 0.001$) of either low TA:DIC (< 1.16) or high TA:DIC (> 1.16) in samples collected from the photic zone. 16S ASVs are colored by phylum and the top five ASVs with the highest indicator values are labeled. There were two ASVs assigned to SAR11 clade 1a. See table S2 for a full list of indicator ASVs and their mean relative abundance. See Fig. 7 for similar plots of 18S ASVs, profiles of TA:DIC in the photic zone, and the correlation between TA:DIC and in situ pH across transects.

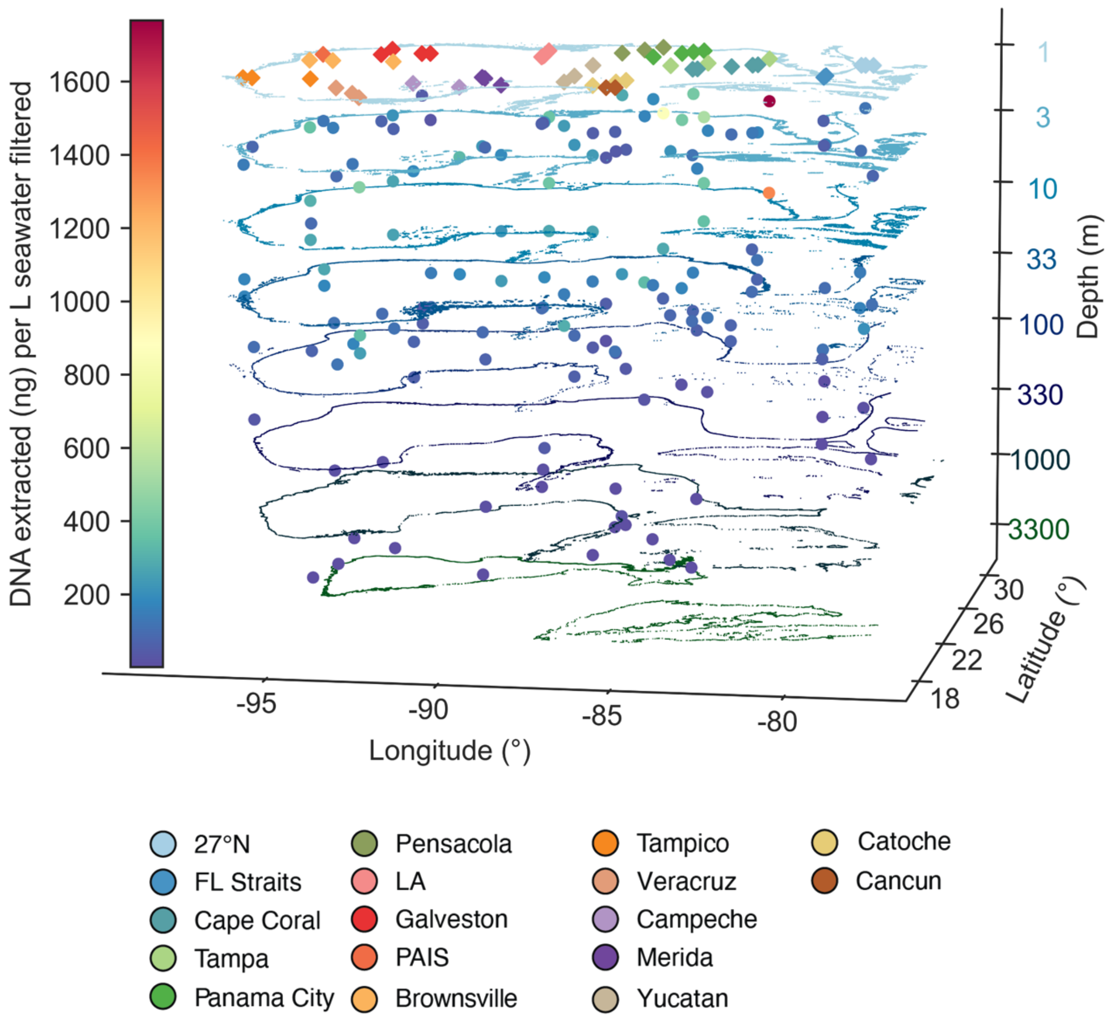


**Figure S13:** Three-dimensional map displaying depth-related (log scale) DNA extracted per volume filtered seawater (ng $l^{-1}$) measured at all sites in the GOM. DNA concentrations were measured following DNA extractions and prior to the first round of PCRs. Shifts in DNA concentration is displayed with a color gradient. Stations are colored by transect at the surface.

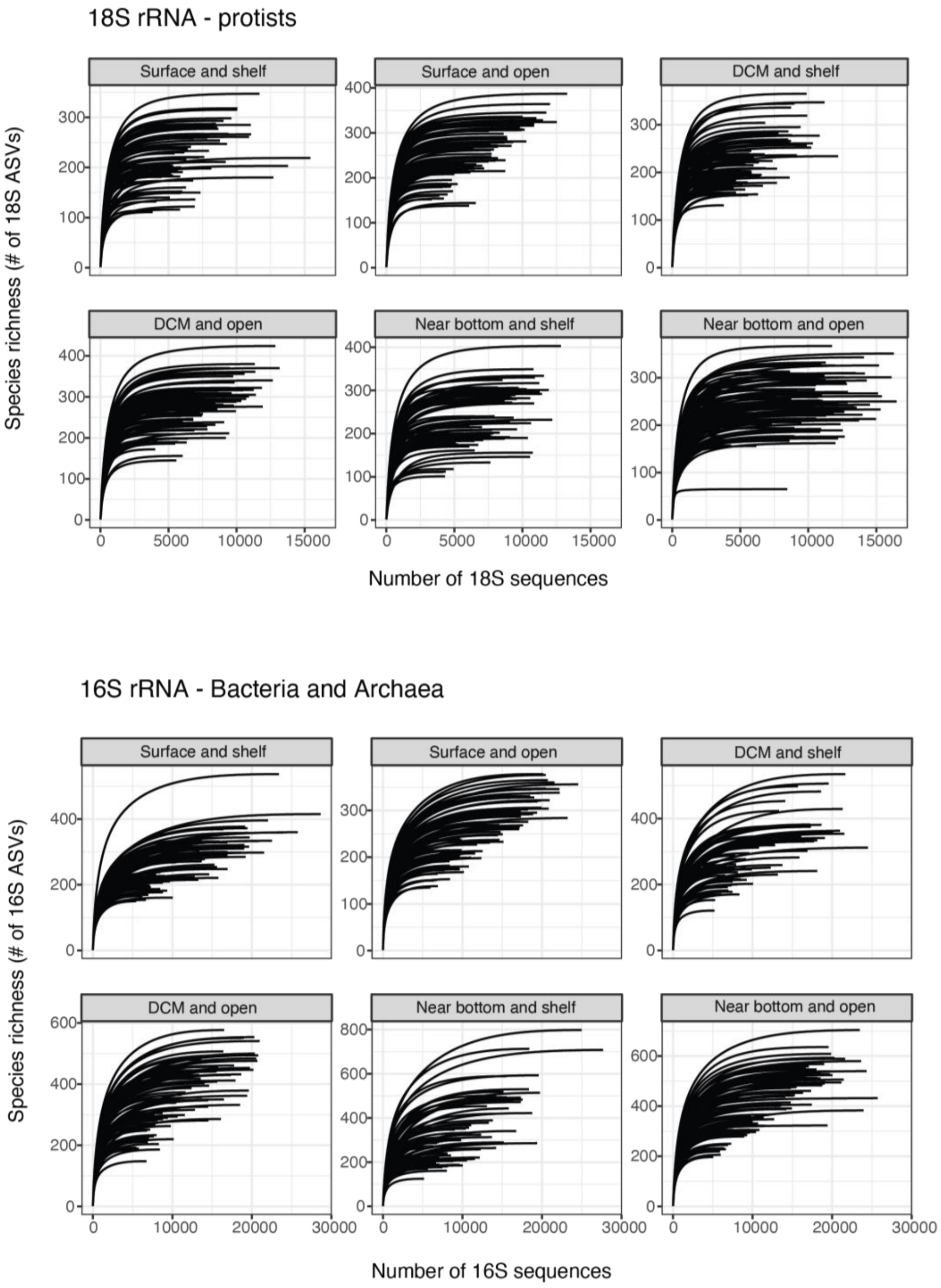


**Figure S14:** Rarefaction curves of 18S (top) and 16S (bottom) species richness (# of ASVs) vs. sequence read counts across all samples and faceted by categorical sampling depth and position of samples on the shelf vs. open ocean. Curves were estimated using a step size of 100.

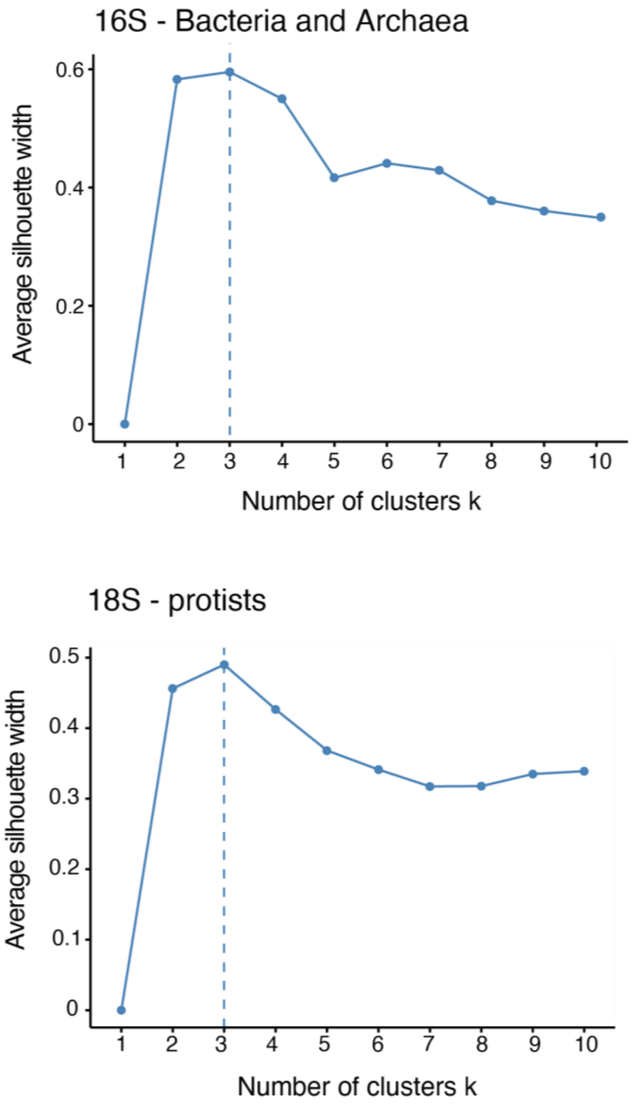


**Figure S15:** Average silhouette widths vs. number of clusters based on hierarchical clustering of 18S (top) or 16S (bottom) Aitchison distance matrices. The optimal number of clusters (three) is indicated by the dotted line and was the same for both amplicon datasets.

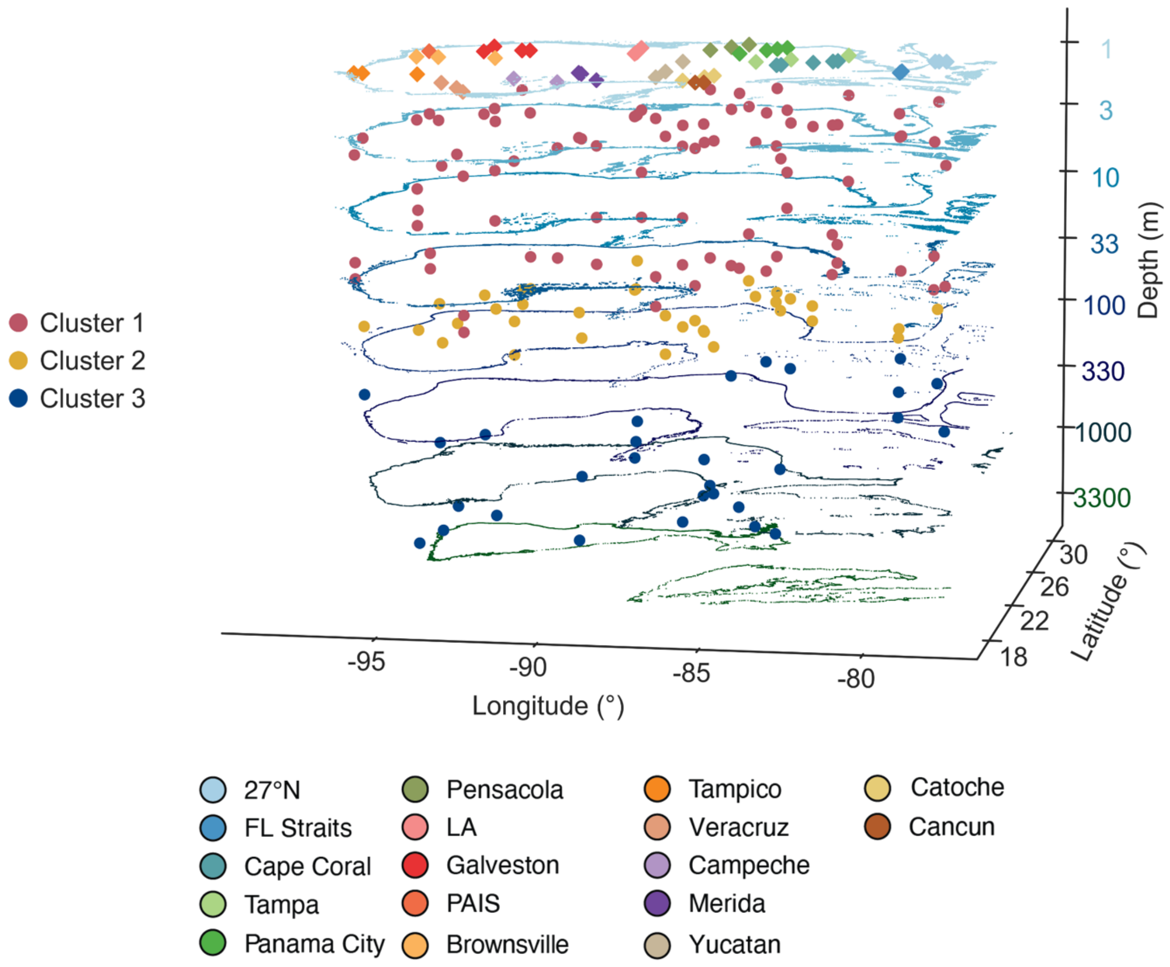


**Figure S16**: Three-dimensional map displaying depth-related position (log scale) of 16S samples collected in the GOM. Stations are colored by transect at the surface, matching Fig. 1A, as well as by their respective clusters (Clusters 1–3). Clusters reflected depth on the shelf and in the open GOM and were similar to those defined for 18S samples (Fig. 1B). All DNA samples were collected in triplicate.

**See Supplementary Files for Tables S1 to S3**

**Table S1:** Spearman correlations between environmental factors for 16S and 18S datasets in Clusters 1–3 (separate sheets). pH values were recalculated based on in situ conditions. TA = total alkalinity and DIC = dissolved inorganic carbon. Variables were selected for Cluster 1 models based on low collinearity (Spearman $r_S < 0.7$ or $> -0.7$) that resulted in a variance inflation factor (VIF) < 10.

**Table S2:** Summary results of indicator analysis for 16S and 18S samples (separate sheets) based on high (> 1.16) or low (< 1.16) TA:DIC ratios in the photic zone. Significant indicator values are shown at the ASV level, along with taxonomic information for each 16S (phylum, genus, and species) and 18S ASV (division, genus, and species). Taxonomic levels were assigned via the PR2 and SILVA databases for 18S and 16S samples, respectively. Average relative abundance (%) of each ASV in the photic zone is also included. Prokaryotic ASVs are labeled as "bASV" to denote them from protists.

**Table S3:** Environmental parameters measured at each site and depth that was sampled for DNA on GOMECC-4 and included in data analysis. Samples are distinguished by transect (region), station number, categorial depth (surface, DCM, and near bottom), and replicate (A–C). Distance to shore indicates position of samples on the continental shelf (inshore; < 200 m) vs. in the open GOM (offshore; > 200 m). Volume filtered for discrete DNA samples is included, which was used to estimate DNA yield (ng $l^{-1}$). The total alkalinity:dissolved inorganic carbon (TA:DIC) ratio was calculated and used for indicator analysis based on manual categories of low (< 1.16) vs. high TA:DIC (> 1.16). Cluster designations are also provided (Clusters 1–3) for 16S and 18S samples.
